# Vav2 is a master regulator of repair against bacterial pore-forming toxins

**DOI:** 10.1101/2025.07.31.668026

**Authors:** Victor Gbenga Kayejo, Ashlee Hensley, Tejal Katore, Peter A. Keyel

## Abstract

Despite antibiotic therapy, 25-35% of patients with necrotizing soft tissue infections (NSTIs) die. The etiologic agents for NSTIs include *Streptococcus pyogenes* and *Clostridium perfringens*, which secrete the cholesterol-dependent cytolysins (CDCs) streptolysin O and perfringolysin O to disrupt cell membranes. While cells resist this damage by activating Ca^2+^-dependent repair pathways, including MEK-dependent microvesicle shedding, dysferlin-mediated patch repair, and annexin-mediated membrane clogging, the upstream regulators of these responses have remained elusive. Here, we demonstrated that the Rac GEF Vav2 accounts for almost all of Ca^2+^-dependent repair against CDCs. Inhibiting or knocking down Vav2 sensitized multiple cell types to CDCs, whereas blocking other Rac GEFs did not. Mechanistically, Vav2 triggered the critical MLK3-MEK-dependent repair pathway. MEK activation rescued repair in Vav2-inhibited cells. Blocking dysferlin or annexins failed to increase damage beyond Vav2 inhibition, suggesting Vav2 coordinates multiple repair pathways. Thus, Vav2 controls multiple Ca^2+^-activated repair pathways that protect cells from CDCs produced during NSTIs.

**Teaser:** This one protein accounts for almost all Ca2+-dependent repair by activating at least 3 distinct pathways.

## INTRODUCTION

Severe bacterial infections that destroy skin, muscle, and soft tissue kill 20-40% of patients, even with medical care (Hakkarainen *et al*, 2014; Jabbour *et al*, 2016). These necrotizing soft tissue infections (NSTIs) are caused by bacteria such as *Streptococcus pyogenes* and *Clostridium perfringens* (Thapa *et al*, 2020). The virulence of these bacteria comes in part from the secreted pore-forming toxins (PFTs) streptolysin O (SLO) and perfringolysin O (PFO) (Awad *et al*, 2001; Limbago *et al*, 2000). SLO and PFO are cholesterol-dependent cytolysins (CDCs), so they bind to cholesterol and create pores in the cell membrane, thereby causing membrane damage (Thapa *et al*., 2020). Damage to epithelial, endothelial, and immune cells enhances bacterial spread into tissues and evasion of the immune response (Los *et al*, 2013; Thapa *et al*., 2020). Failure to resolve membrane damage leads to cell death, so it is crucial to understand how cells repair membrane damage.

Membrane repair describes a collection of potentially interrelated biological processes that reseal the membrane. Resealing mechanisms can vary by cell type, source of damage, and extent of damage, which complicates pathway determinations (Cooper & McNeil, 2015; Jimenez & Perez, 2017). In general, cells sense membrane damage by ion flux, especially Ca^2+^ (Cooper & McNeil, 2015; Jimenez & Perez, 2017). Upon damage, three Ca^2+^-activated repair mechanisms reseal the membrane by clogging the breach, patching the breach, and/or sequestering and eliminating the breach by shedding damaged membranes as microvesicles (Thapa *et al*., 2020). Clogging is mediated by annexins and transglutaminases, which form crystalline lattices and plugs to block the breach (Demonbreun *et al*, 2016b; Jimenez & Perez, 2017; Roostalu & Strahle, 2012). Patch repair is the hetero- and homotypic fusion of vesicles with the membrane, driven by C2 domain-containing proteins, including synaptotagmins, copines, and the muscle-specific protein dysferlin (Alves *et al*, 2022; Chakrabarti *et al*, 2003; McNeil & Kirchhausen, 2005). Microvesicle shedding is the sequestration and shedding of pore-forming toxins on microvesicles, controlled by lipids (Keyel *et al*, 2011), Endosomal Sorting Complex Required for Transport (ESCRT) recruitment (Jimenez *et al*, 2014), and MEK activation (Ray *et al*, 2022). Elevation of ceramide and other proteins can protect cells from damage (Haram *et al*, 2023; Ray *et al*., 2022; Schoenauer *et al*, 2019) via unknown mechanisms. While previously thought to be a repair mechanism, we and others showed that endocytosis occurs after repair to clean up the damage (Keyel *et al*., 2011; Nygard Skalman *et al*, 2018; Romero *et al*, 2017; Schoenauer *et al*., 2019; Stefl *et al*, 2024; Thapa & Keyel, 2023). The relative weights of these repair pathways vary by toxin type because patch repair is more important to resist the small pore-forming toxin aerolysin, whereas microvesicle shedding is more critical to stop CDCs (Thapa & Keyel, 2023). The signaling pathways coordinating these pathways downstream of Ca^2+^ influx are unknown.

Since many repair proteins contain Ca^2+^-binding domains, one hypothesis is that Ca^2+^ influx is sufficient to drive repair. For example, dysferlin, annexins, and copines all contain Ca^2+^-binding domains and are recruited to damage sites (Alves *et al*., 2022; Boye *et al*, 2017; Demonbreun *et al*, 2016a; Kayejo *et al*, 2023). However, live cell imaging shows that cells are flooded with Ca^2+^ throughout the cell upon damage (Babiychuk *et al*, 2009a, 2011; Thapa & Keyel, 2023). With elevated Ca^2+^ throughout the cell, it is unclear how Ca^2+^-sensitive proteins are targeted specifically to the site of damage. Therefore, additional signaling pathways are needed to coordinate the repair response.

Our recent work showed that an atypical MAP kinase signaling pathway involving Mixed Lineage Kinase 3 (MLK3) and Mitogen-Activated Protein Kinase Kinase (MEK), but not Extracellular Regulated Kinase (ERK), accounts for 70% of Ca^2+^-dependent repair. This signaling axis triggers annexin A2 membrane translocation and microvesicle shedding (Ray *et al*., 2022). When this pathway is blocked, annexin A1 and annexin A6 try to compensate by translocating to the membrane faster (Ray *et al*., 2022). However, the upstream signaling proteins that activate MLK3 remain unknown.

One potential upstream signaling partner active during membrane repair is the Rho GTPase family. Rho GTPases such as Rho, Rac, and cdc42 are each implicated as signaling proteins that promote actin remodeling and wound healing after damage in *Xenopus* oocytes (Benink & Bement, 2005; Moe *et al*, 2021), and mammalian cells (Iliev *et al*, 2007; Lam *et al*, 2018). These small GTPases have many downstream interaction partners, positioning them to execute cell signaling programs in response to extracellular stimuli like damage. Notably, both cdc42 and Rac interact with MLK3 via the latter’s cdc42/Rac interactive binding motif (CRIB motif) (Du *et al*, 2005). Activation of these GTPases is further controlled by guanine nucleotide exchange factors (GEFs). For example, Rac1 can be activated by Vav proteins, T-lymphoma invasion and metastasis-inducing protein-1 (Tiam1), Trio, DOCK1/2, Dbl’s big sister (Dbs), or Faciogenital dysplasia 5 (Fgd5) (Maldonado *et al*, 2020). Thus, specific GEFs could activate Rho GTPases, which trigger MLK3-MEK-dependent membrane repair pathways in response to bacterial CDCs.

Here, we tested the hypothesis that the Rac GEF Vav activates MLK3-MEK repair of membrane damage caused by bacterial CDCs. Blocking Vav2, but not other Rac GEFs, nor cdc42 and Rho, halted the MLK3/MEK signaling axis. Moreover, Vav activation accounted for almost all Ca^2+^-dependent repair against CDCs in multiple cell lines. We propose that this additional repair occurs because Vav2 acts upstream of dysferlin and annexins A1, and A2. Our results suggest that Vav2 is an upstream master regulator coordinating multiple membrane repair pathways to protect against CDCs.

## RESULTS

### Rac inhibition sensitizes cells to bacterial pore-forming toxins

To determine the contribution of Rho GTPases to repair, we blocked each individually or in combination. We first validated the functional activity of each inhibitor by measuring actin cytoskeleton disruption in HeLa cells treated with each inhibitor. Cdc42, Rho, and Rac inhibitors disrupted actin fiber organization (Supplementary Fig S1A, B). These data demonstrate that the inhibitors were active. Next, we titrated the inhibitors to determine at what dose, if any, they altered cell sensitivity to CDCs. We pretreated HeLa cells with 5- 20 μM of Rac inhibitor EHop016, cdc42 inhibitor ML141, Rho inhibitor Y16 or ROCK inhibitor Y27632, then challenged cells with SLO or PFO. To measure toxin activity, we used hemolytic units instead of mass because it normalizes toxin activity across toxin preparations and experiments (Supplemental Table S1). We fit dose-response curves to a logistic model to determine the lethal concentration of 50% (LC_50_), the toxin dose needed to kill 50% of the cells (Haram *et al*, 2022), which enables us to compare the impact of each inhibitor. Inhibitors were not toxic to cells unchallenged with CDCs. When challenged with SLO or PFO, Rac inhibition increased the sensitivity of HeLa cells to bacterial toxins by 80-90% compared to vehicle-treated (control) cells (Fig 1A, Supplementary Fig S1C). In contrast, cdc42 inhibition did not alter cell sensitivity to either toxin compared to DMSO-treated control cells (Fig 1B Supplementary Fig S1D).

**Figure 1.**
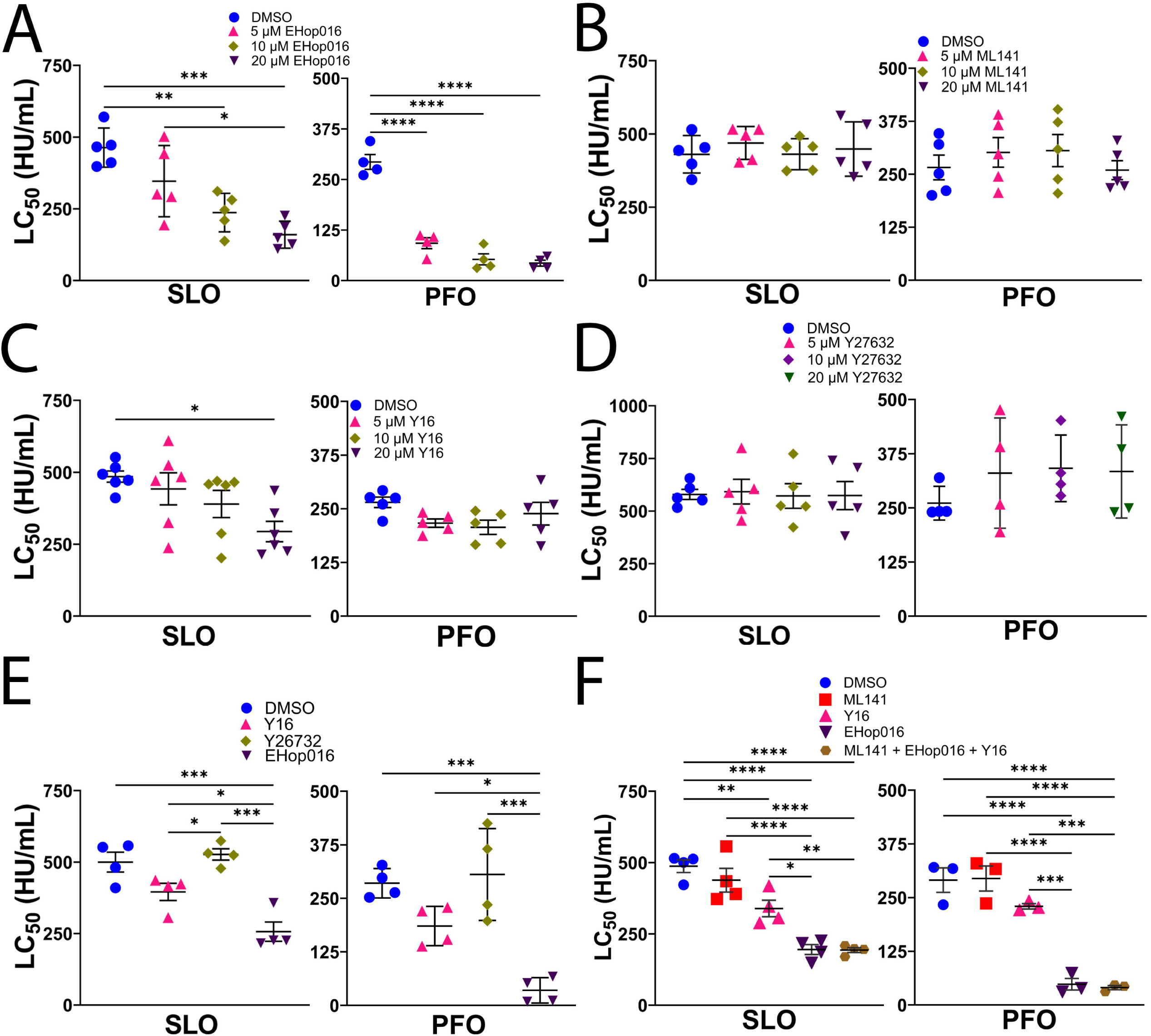
Blocking Rac activation sensitizes cells to bacterial pore-forming toxins. HeLa cells were serum-starved for 30 min, treated with DMSO or 5, 10, or 20 μM **(A)** Rac inhibitor EHop016, (B) cdc42 inhibitor ML141, (C) Rho inhibitor Y16, (D) ROCK inhibitor Y27632, (E) 20 μM Y16, Y27632, or EHop016 or (F) 20 μM ML141, EHop016, Y16, or all three, and challenged with 31-2000 HU/mL SLO or PFO for 30 min at 37°C. Propidium iodide (PI) uptake was analyzed by flow cytometry. The LC_50_ was calculated as described in the methods. Graphs show independent experiments and the mean ±S.E.M, n=5 (A, B, D (SLO), B, C, (PFO)), n=4 (A, D, E, (PFO), D, E, F (SLO)), n=6 (C, SLO), or n=3 (F, PFO). *p<0.05, **p<0.01, ***p<0.001, ****p<0.0001 denote statistical significance using repeated-measures ANOVA between groups with Tukey post-test.

Similarly, Rho inhibition increased sensitivity to SLO at 20 µM but not to PFO (Fig 1C, Supplementary Fig S1E). Since Rho can signal through ROCK1/2, we tested if ROCK1/2 signaling is involved in SLO sensitivity. ROCK1/2 inhibition did not alter cell sensitivity to either toxin (Fig 1D, Supplementary Fig S1F). When we compared Rho, Rac, and ROCK1/2 inhibition, the increase in sensitivity to SLO and PFO of Rac-inhibited cells was statistically significant compared to vehicle and other inhibitors (Fig 1E, Supplementary Fig S1G). We then combined inhibitors to determine any redundancies in CDC sensitivity. We found no increase in sensitivity when other inhibitors were combined with Rac inhibition compared to Rac inhibition alone (Fig 1F, Supplementary Fig S1H). Therefore, we conclude that the Rac inhibitor EHop016 sensitizes cells to SLO and PFO.

Since Rac inhibition sensitizes cells to SLO and PFO, we next determined if Rac was needed for repair against other toxins. The small pore-forming toxin aerolysin is resisted primarily by patch repair instead of microvesicle shedding (Thapa & Keyel, 2023). We challenged Rho, ROCK1/2, or Rac-inhibited HeLa cells with aerolysin. There was a trend towards increased HeLa cell sensitivity to aerolysin upon Rac inhibition (Supplementary Fig S1I). Interestingly, cdc42 inhibition increased cellular resistance to aerolysin challenge (Supplementary Fig S1I). Overall, these data suggest that Rac triggers specific repair pathways to protect cells from CDCs.

To validate the importance of Rac for repair, we examined multiple Rac inhibitors and multiple cell types. We treated HeLa cells with a second Rac inhibitor, EHT1864, before SLO or PFO challenge. We found increased cell sensitivity, similar to EHop016 (Fig 2A, Supplementary Fig S2A). Next, we evaluated the role of Rac inhibition in membrane repair in other cell types. We inhibited Rac using EHT1864 in primary bone marrow-derived macrophages (BMDMs) from C57BL/6 (B6) mice or using EHop16 in HEK cells and C2C12 murine myoblasts prior to CDC challenge. Rac inhibition increased cell death compared to control cells in all cell types (Fig 2B-D, Supplementary Fig S2B-D). Therefore, we conclude that multiple mammalian cell types rely on this pathway for repair.

**Figure 2.**
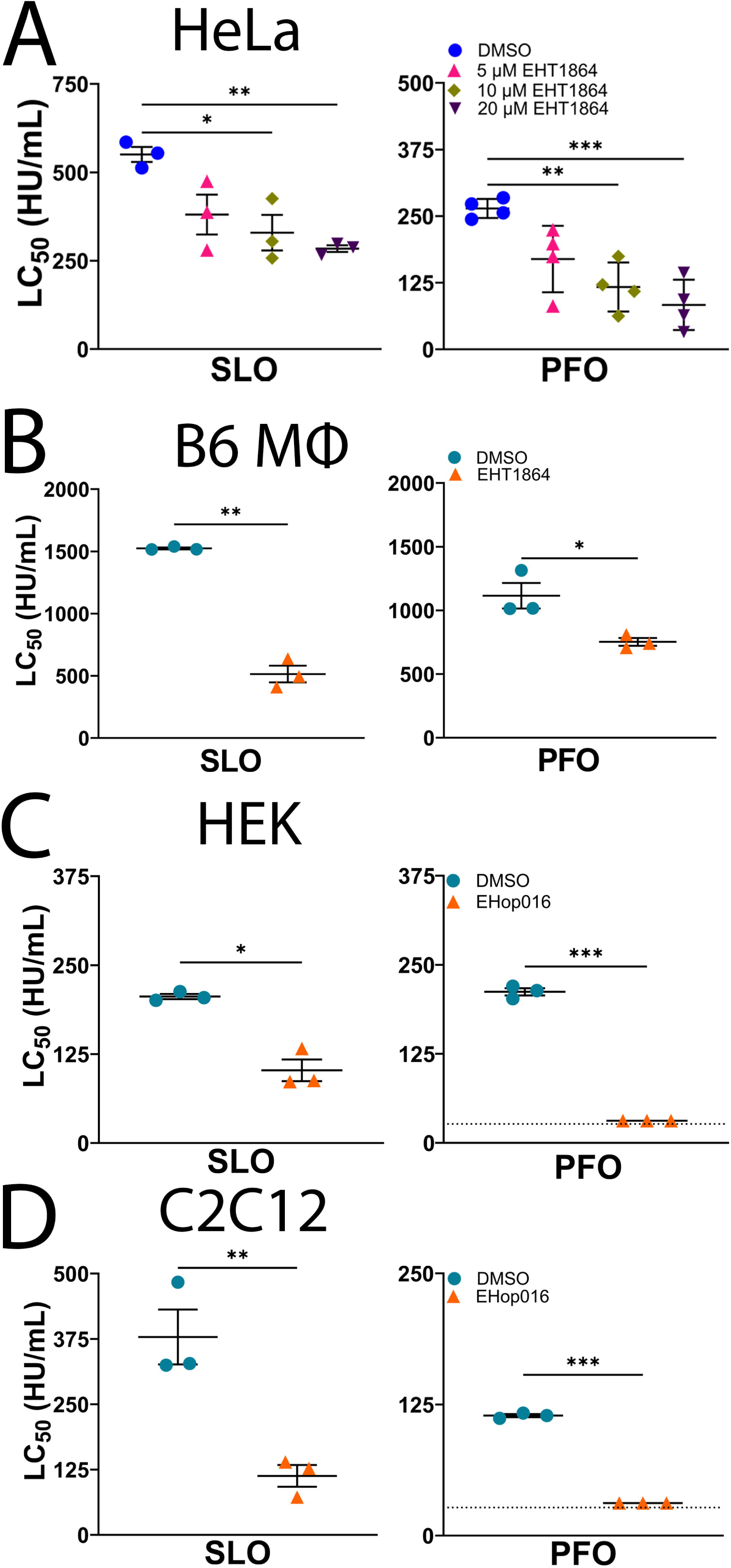
Multiple cell types are susceptible to toxins upon Rac inhibition. (A) HeLa, (B) bone marrow-derived macrophages (B6 M), (C) HEK, or (D) C2C12 cells were serum-starved for 30 min, treated with (A) DMSO or 5, 10, or 20 μM Rac inhibitor EHT1864, DMSO or (B) 20 μM EHT1864, or (C, D) 20 μM EHop016 for 30 min, challenged with 31-4000 HU/mL SLO or PFO at 37°C for 30 min, and analyzed by flow cytometry for PI uptake. The LC_50_ was calculated as described in the methods. The dotted line indicates the limit of detection. Points on this line had LC_50_<15 HU/ml (C, D, PFO). Graphs display independent experiments and the mean ±S.E.M for n=3 (A-D) or n=4 (A, PFO). *p<0.05, **p<0.01, ***p<0.001 denote statistical significance using repeated-measures ANOVA between groups with Tukey post-test.

### Vav mediates toxin resistance

We next narrowed the number of Rac GEFs that could be responsible for Rac activation during membrane repair. While EHT1864 blocks all Rac GEFs, including Vav (Shutes *et al*, 2007), some Rac inhibitors only block a subset of Rac GEFs. For example, EHop016 inhibits Vav activation of Rac (Montalvo-Ortiz *et al*, 2012). To determine if other GEFs promote Rac-dependent repair, we blocked Tiam1 and Trio using the inhibitor NSC23766 (Gao *et al*, 2004). We validated the functional activity of NSC23766 by checking for actin cytoskeleton disruption (Supplementary Fig S3A). In contrast to Rac inhibition via EHop016, inhibition with NSC23766 failed to increase the sensitivity of HeLa cells to toxin challenge (Fig 3A, Supplementary Fig S3B). This suggests that Tiam1 and Trio are not involved in activating Rac during repair, but Vav is.

**Figure 3.**
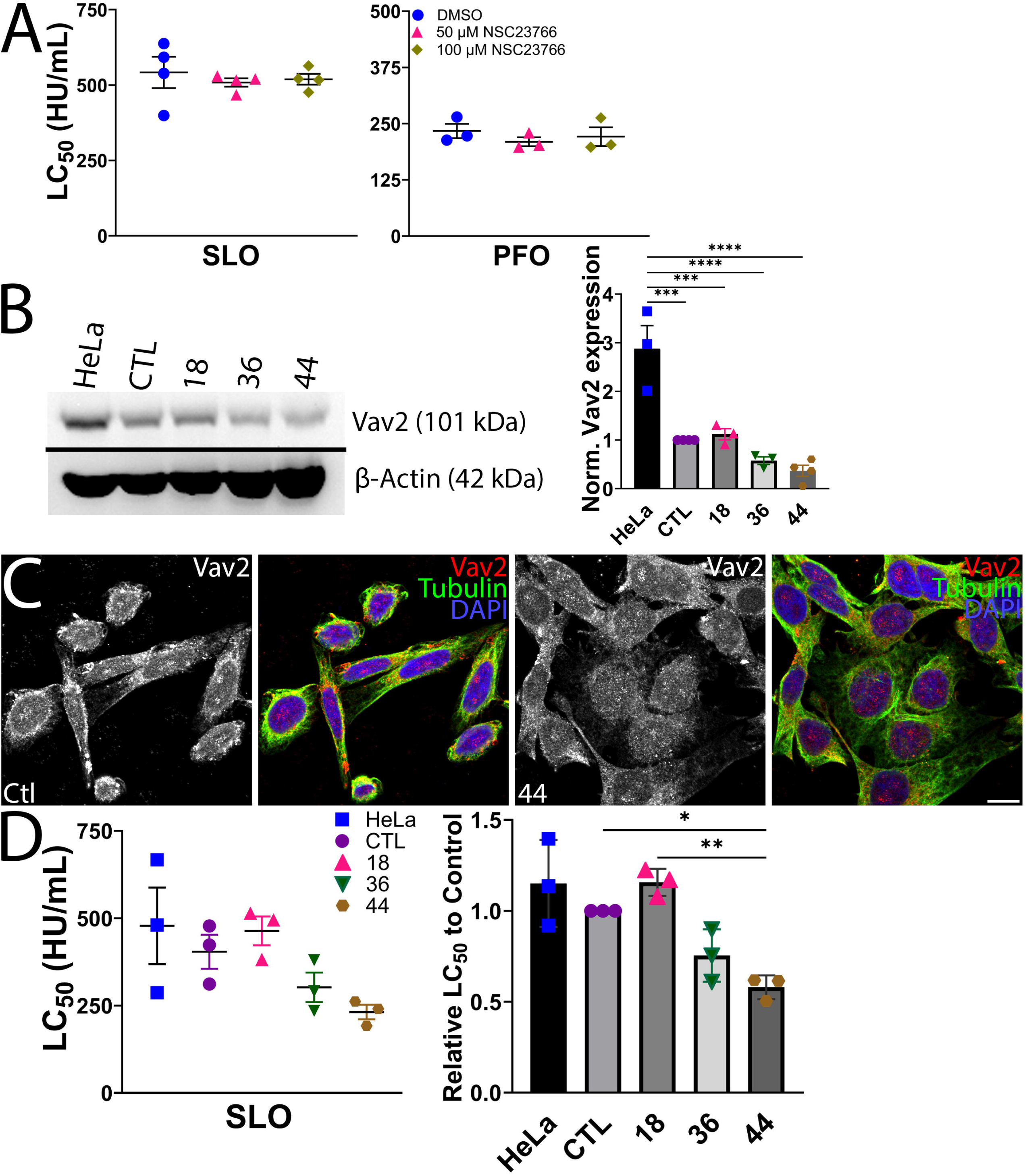
Vav2 promotes membrane repair. (A) HeLa cells were serum-starved for 30 min, treated with DMSO or 50 or 100 μM NSC23766 for 5, 10, or 20 μM for 30 min, then challenged with 31-2000 HU/mL SLO or PFO for 30 min at 37°C. PI uptake was analyzed by flow cytometry. The LC_50_ was calculated as described in the methods. (B-D). HeLa cells were transfected with negative control guide RNA or guide RNA targeting Vav2. Cells were harvested, single cell cloned and expanded. (B) Western blot of untransfected HeLa cells, one CRISPR negative control clone (CTL), and three Vav2 clones (18, 36, and 44). Western blots from multiple passages were quantitated and relative Vav2 expression determined. (C) CTL or Vav2 CRISPR clone 44 were fixed, permeabilized and stained with anti-Vav2 (red), anti-tubulin (green) and DAPI (blue). (D) Cells were challenged with 31-2000 HU/mL SLO for 30 min at 37°C. PI uptake was analyzed by flow cytometry. The LC_50_ relative to CTL cells was determined for each experiment. Graphs show independent experiments, and the mean ±S.E.M for n=4 (A, SLO) or n=3 (A PFO, D) independent experiments. The blot and micrographs represent three independent experiments. *p<0.05, **p<0.01 denote statistical significance using repeated-measures ANOVA between groups with Tukey post-test.

HeLa cells express Vav2 and Vav3 because Vav1 is restricted to hematopoetic cells. Since Vav3 has regulatory roles (Bustelo, 2014), we targeted Vav2 gene expression using CRISPR. Following two attempts at CRISPR, we grew out clones with Vav2 expression reduced ∼67% compared to CRISPR control cells, which was stable across passages (Fig 3B). We examined the heterogeneity of our cells to determine if the residual Vav2 expression was due to mosaic Vav2 expression, or if all cells lost two of the three Vav2 alleles present in HeLa cells (Landry *et al*, 2013). We found Vav2 CRISPR cells showed even, residual Vav2 staining (Fig 3C). If the Vav2 cells had mosaic expression, the variance in Vav2 intensity is expected to be distinct from the variance present in control cells. We tested this hypothesis using Levene’s test. None of our three replicates showed unequal variance between control and Vav2 CRISPR cells. This suggests we eliminated two of the three Vav2 alleles, instead of having mosaic expression. Cell sensitivity to SLO increased when Vav2 was >50% reduced in cells (Fig 3D, Supplementary Fig S3C). Based on these data, we conclude that Vav2 drives membrane repair.

Since Vav2 is a phosphoprotein, we next tested several upstream kinases that could activate Vav2 during membrane repair. We inhibited Src/Syk tyrosine kinases using the inhibitor 3,4-methylenedioxy-β-nitrostyrene (MNS) (Wang *et al*, 2006), and focal adhesion kinase and Pyk2B with the inhibitor PF431396 (Han *et al*, 2009). However, neither inhibitor increased CDC sensitivity (Fig 4A, B, Supplementary Fig S3D, E). Similarly, siRNA knockdown of Pyk2B failed to increase HeLa cell sensitivity compared to control cells (Fig 4C, D, Supplementary Fig S3F). These results suggest that Vav2 promotes membrane repair independent of the tyrosine kinases Src, Syk, focal adhesion kinase, and Pyk2B.

**Figure 4.**
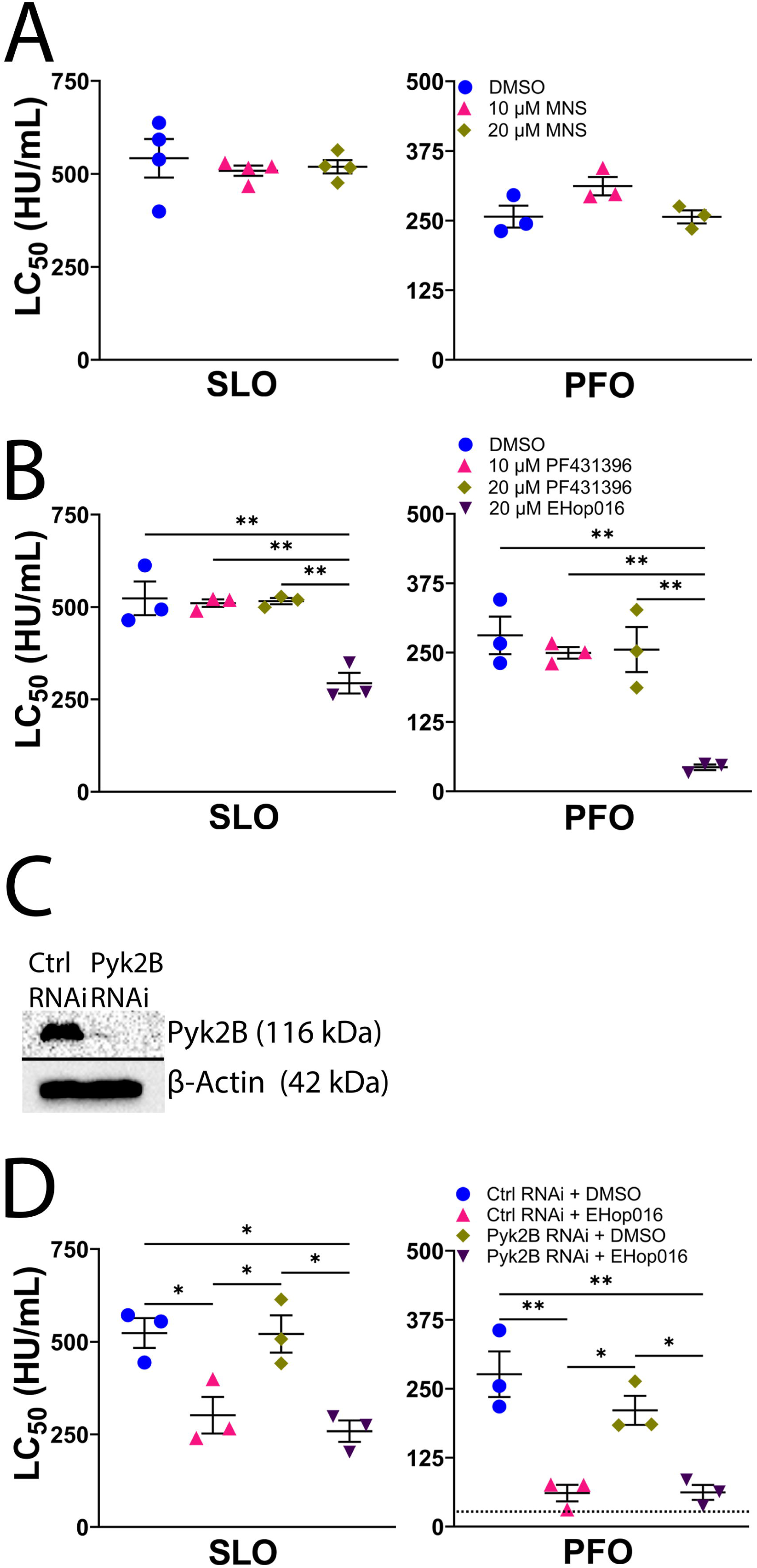
Vav2 promotes membrane repair independent of the tyrosine kinases Src, Syk, focal adhesion kinase, and Pyk2. HeLa cells were serum-starved for 30 min, treated with DMSO or 10 or 20 μM (A) Src/Syk inhibitor MNS or (B) FAK/Pyk2 inhibitor PF431396 or EHop016 for 30 min, and then challenged with 31-2000 HU/mL SLO or PFO for 30 min. HeLa cells were transfected with control (Ctrl) or Pyk2-siRNA for 72 hours, and (C) lysed for western blot analysis or (D) serum-starved for 30 min, treated with DMSO or 20 μM EHop016, and challenged with 31-2000 HU/mL SLO or PFO for 30 min. PI uptake was analyzed by flow cytometry. The LC_50_ was calculated as described in the methods. Graphs show independent experiments, and the mean ±S.E.M for n=4 (A, SLO) or n=3 (A PFO, B, D) independent experiments. The blot represents three independent experiments. *p<0.05, **p<0.01 denote statistical significance using repeated-measures ANOVA between groups with Tukey post-test.

### Vav2 promotes Ca^2+^-dependent repair upstream of MLK3-MEK signaling

To determine if Vav-mediated membrane repair is activated by Ca^2+^ influx, we pretreated HeLa cells with EHop016, with or without Ca^2+^-chelation using EGTA. EGTA alone increased cell sensitivity, consistent with prior results (Cooper & McNeil, 2015; Keyel *et al*., 2011; Ray *et al*., 2022; Romero *et al*., 2017; Thapa & Keyel, 2023) (Fig 5A, Supplementary Fig S4A). When EGTA was combined with Vav inhibition, no additive effects were observed, suggesting that Vav acts in the Ca^2+^-dependent pathways (Fig 5A, Supplementary Fig S4A). Thus, Vav-dependent repair is activated after Ca^2+^ influx.

**Figure 5.**
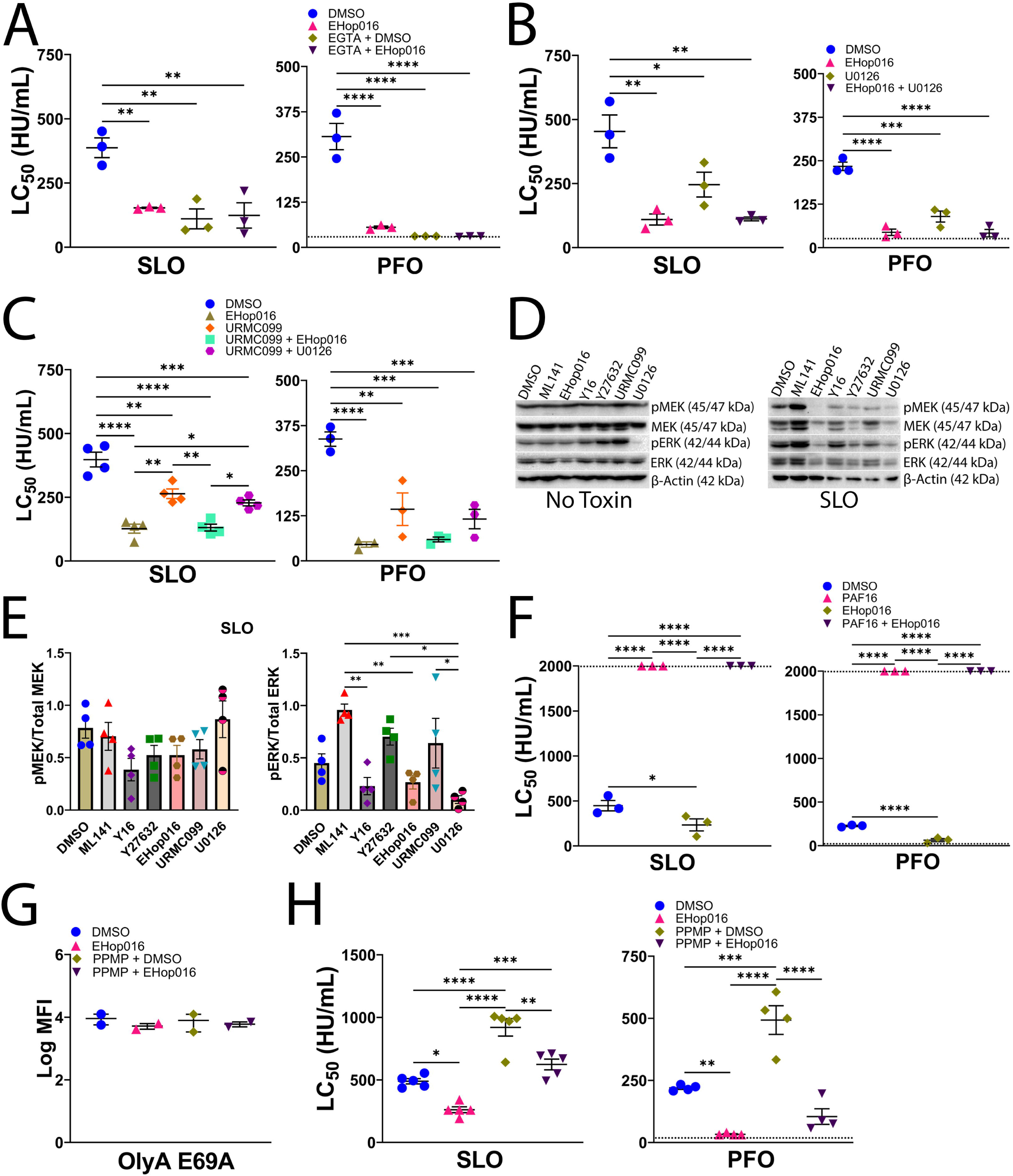
Vav2 promotes Ca^2+^-dependent repair upstream of MLK3-MEK signaling and parallel to ceramide-mediated repair. (A-C) HeLa cells were serum-starved for 30 min and pretreated with DMSO and/or a combination of: 20 μM Rac inhibitor EHop016, calcium chelator EGTA, 20 μM MEK inhibitor U0126, or 20 μM MLK3 inhibitor URMC099 and challenged with 15-2000 HU/mL SLO or PFO for 30 min at 37°C. PI uptake was analyzed by flow cytometry. The LC_50_ was calculated as described in the methods. Where indicated, 2 mM EGTA was used instead of 2 mM CaCl_2_. (D, E) HeLa cells treated with DMSO or 20 μM ML141, Y16, Y27632, EHop016, or URMC099 were challenged with nothing or sub-lytic SLO (250 HU/mL) for 30 min, and analyzed by western blot for the indicated antibodies. The ratio of phospho proteins to total protein was calculated. (F). HeLa cells were treated and analyzed as in (A) using 20 μM EHop016 and/or MEK activator PAF16. (G, H). HeLa cells were treated with 20 μM PPMP for 72 h at 37°C. Then, cells were serum-starved for 30 min at 37°C, treated with DMSO or 20 μM EHop016 for 30 min, and challenged with either (G) 20 μg/mL Ostreolysin A (OlyA) E69A–mCherry, for 5 min, or (H) 31-2000 HU/mL SLO or PFO for 30 min. Graphs display independent experiments and the mean ±S.E.M for n=2 (G), n=3 (A, B, F, (SLO), A, B, C, F, (PFO)) or n=4 (C, E, (SLO), E, H, (PFO)) or n=5 (H, SLO). The western blot represents 4 independent experiments. The dotted line indicates the limit of detection. Points on this line had LC_50_<15 HU/ml (A, (PFO)), LC_50_<31 HU/ml (B, (PFO)), or LC_50_>2000 HU/ml (F). *p<0.05, **p<0.01, ***p<0.001, ****p<0.0001 denote statistical significance using repeated-measures ANOVA between groups with Tukey post-test.

We next determined if Vav2 acts upstream of MLK3-MEK signaling. We tested this hypothesis by blocking MEK with the previously characterized MEK inhibitor U0126 (Ray *et al*., 2022), and/or Vav activity with EHop016 in HeLa cells before toxin challenge. After toxin challenge, both inhibitors reduced cell survival (Fig 5B, Supplementary Fig S4B). However, inhibition of both Vav and MEK did not further exacerbate toxin-induced cell death compared to Vav inhibition alone (Fig 5B, Supplementary Fig S4B). These data suggest that Vav and MEK are in the same repair pathway. Next, we blocked Vav activity and/or MLK3 and compared the inhibition to that achieved by blocking both MLK3 and MEK. Upon toxin challenge, Vav inhibition blocked more repair to SLO and PFO than MLK3 inhibition alone or MLK3-MEK inhibition combined (Fig 5C, Supplementary Fig S4C). These data suggest that Vav acts upstream of the MLK3-MEK signaling pathway during membrane repair.

To confirm that Vav acts upstream of MLK3 and MEK, we measured the phosphorylation of MEK and ERK after Vav inhibition. While repair is ERK-independent (Ray *et al*., 2022), ERK phosphorylation is a readout for MEK activity. Relative to control cells, at baseline, ERK and MEK phosphorylation were unchanged for all inhibitors except MEK blockade, consistent with the ability of Raf and other kinases to activate MEK and ERK (Fig 5D, E). However, upon challenge with a sub-lytic dose of SLO, inhibition of Rho, Rac, or MEK all decreased ERK phosphorylation, whereas cdc42 inhibition increased ERK phosphorylation (Fig 5D, E). If MEK acts downstream of Vav during repair, we predict forced MEK activation would override Vav inhibition and protect cells from CDCs. We tested this hypothesis by pretreating HeLa cells with EHop016 and/or the MEK activator Platelet-activating factor 16 (PAF16) before and during the toxin challenge. As expected, Vav inhibition increased cell sensitivity to toxins (Fig 5F, Supplementary Fig S4D). In contrast, MEK stimulation using PAF16 protected HeLa cells from SLO or PFO, regardless of Vav inhibition (Fig 5F, Supplementary Fig S4D). Based on these data, we conclude that Vav acts upstream of MLK3 and MEK signaling during membrane repair.

### Vav and ceramide-mediated repair are parallel

Since Vav acts upstream of MLK3 and confers greater protection to cells, we tested other Ca^2+^-dependent repair pathways to determine if they act in the same pathway as Vav or a parallel pathway to Vav. Ceramide contributes to repair (Haram *et al*., 2023; Ray *et al*., 2022; Schoenauer *et al*., 2019), via a parallel pathway to MEK (Ray *et al*., 2022). We tested the hypothesis that Vav is involved in ceramide-driven repair. To elevate ceramide without sphingomyelin depletion, we used d-threo-1-phenyl-2-hexadecanoylamino-3-morpholino-1-propanol ·HCl (PPMP) for 72 hours to induce accumulation of ceramide in cells (Dany & Ogretmen, 2015). Neither PPMP treatment nor Vav inhibition altered total surface sphingomyelin levels (Fig 5G). When HeLa cells were challenged with toxins, PPMP protected cells (Fig 5H, Supplementary Fig S4E), as previously observed (Ray *et al*., 2022). While EHop016 reduced the LC_50_ of PPMP-treated cells, it failed to make cells as sensitive as Vav inhibition alone (Fig 5H, Supplementary Fig S4E).

Since Vav inhibition partly overcame ceramide protection, we conclude that Vav-driven and ceramide-mediated repair are parallel repair pathways.

### Vav may regulate dysferlin and annexin-mediated repair

Next, we investigated the role of Vav in patch repair by determining its role in dysferlin-mediated repair. We inhibited Vav in C2C12 myoblasts stably transfected with control or dysferlin-shRNA before the toxin challenge. Dysferlin knockdown was confirmed by western blot (Fig 6A). Consistent with previous results (Thapa & Keyel, 2023), dysferlin loss increased cell sensitivity to CDCs (Fig 6B, Supplementary Fig S5A). Vav inhibition increased the CDC sensitivity of cells to a similar extent, regardless of dysferlin expression (Fig 6A, B, Supplementary Fig S5A). We interpret these data to suggest that Vav regulates dysferlin-mediated repair to toxin damage.

**Figure 6.**
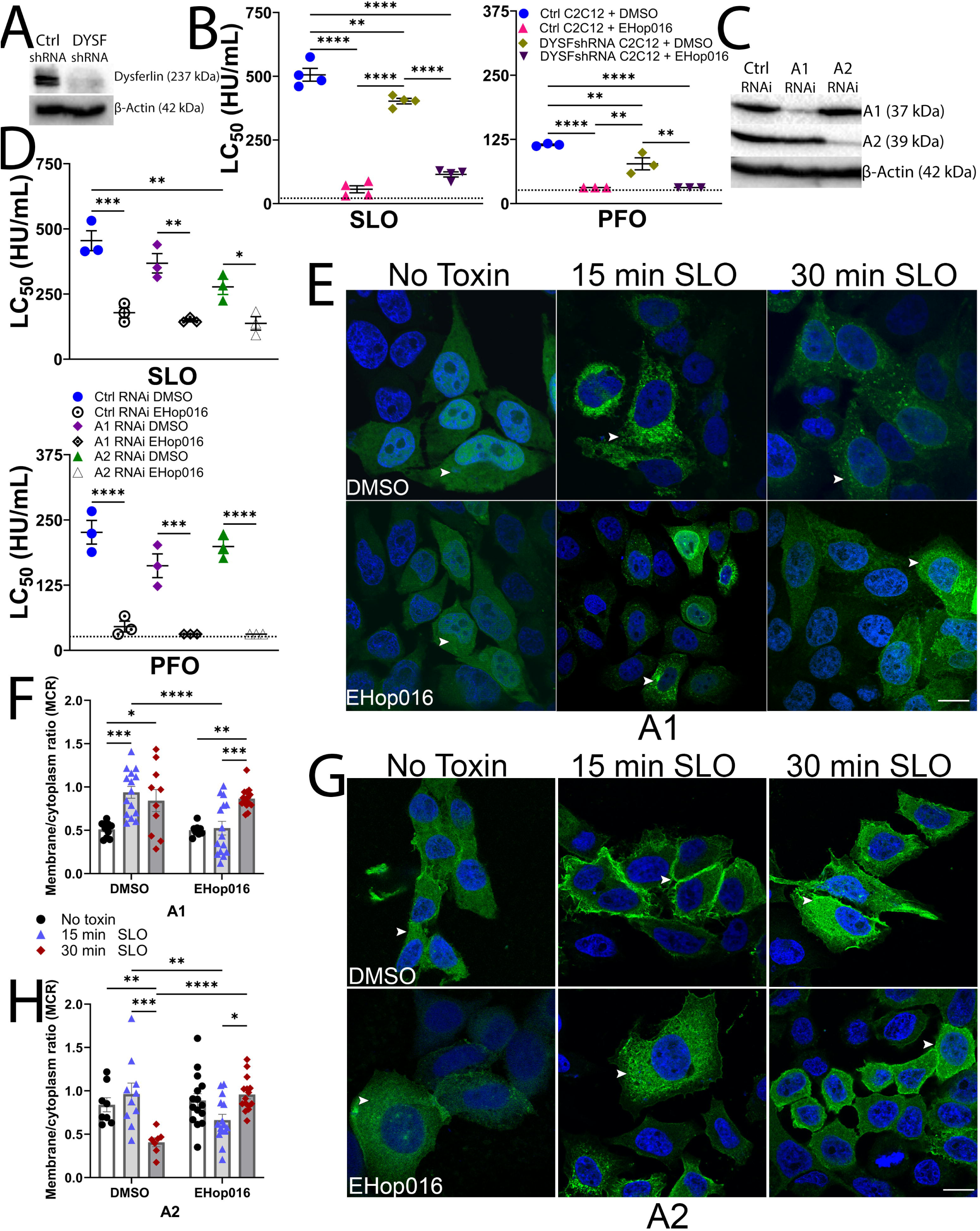
Vav regulates dysferlin and annexin-mediated repair. (A, B). Control shRNA or dysferlin-shRNA C2C12 cells were (A) lysed for western blot analysis, or (B) serum-starved for 30 min, treated with DMSO or 20 μM EHop016, and challenged with 15-2000 HU/mL SLO or PFO for 30 min. PI uptake was analyzed by flow cytometry. (C, D). HeLa cells transfected with control (Ctrl), annexin A1 (A1), or annexin A2 (A2) siRNA for 72 hours were (C) lysed for western blot analysis or (D) serum-starved for 30 min, pre-treated with DMSO or 20 μM EHop016 for 30 min, and challenged with 31-2000 HU/mL SLO or PFO for 30 min. PI uptake was analyzed by flow cytometry. The LC_50_ was calculated as described in the methods. HeLa cells plated on coverslips were transfected with (E, F) A1-YFP or (G, H) A2-GFP, serum-starved for 30 min, pre-treated with DMSO or 20 μM EHop016 for 30 min, and challenged with sub-lytic SLO (250 HU/mL) for 0, 15, or 30 min at 37°C. (E, G) Cells were fixed, stained with DAPI, and imaged. Arrowheads show membrane vs cytoplasmic annexins. (F, H) Translocation of annexins to the membrane was quantified. Graphs show independent experiments and the mean ±S.E.M for n=4 (B, SLO) or n=3 (D (SLO), B, D, (PFO)) or individual cells and the mean ±S.D. from three independent experiments (F, H). The dotted line indicates the limit of detection. Points on this line had LC_50_<15 (B, D (PFO)) or 31 HU/ml (B (SLO)). The blots represent 3 independent experiments. Micrographs represent three independent experiments. Scale bar= 10 μm.*p<0.05, **p<0.01, ***p<0.001, ****p<0.0001 denote statistical significance using repeated-measures ANOVA between groups with Tukey post-test.

Next, we tested the involvement of Vav in annexin-mediated clogging. We depleted annexins A1 (A1), and A2 (A2) by siRNA and confirmed knockdown by western blot (Fig 6C). We then challenged these annexin-depleted cells with SLO or PFO in the presence or absence of Vav inhibition. Consistent with prior results (McNeil *et al*, 2006; Swaggart *et al*, 2014; Thapa & Keyel, 2023), we found that knockdown of each annexin alone increased cell sensitivity to CDCs compared to control cells (Fig 6D, Supplementary Fig S5B). However, A1 or A2 depletion failed to exacerbate cell death due to Vav inhibition after CDC challenge (Fig 6D, Supplementary Fig S5B). These data suggest that Vav acts upstream of multiple annexins during repair against bacterial CDCs.

We next examined the rate of annexin binding to the membrane during Rac inhibition. Our prior results showed that A2 membrane recruitment took over 20 min when MEK was blocked, but A1 had accelerated membrane recruitment (Ray *et al*., 2022). To determine if A1 and A2 both had retarded recruitment or if recruitment was normal, we measured annexin recruitment in vehicle-treated or Vav-inhibited cells at 15 and 30 min post-SLO challenge by high-resolution confocal microscopy. Both annexins were recruited to the membrane from the cytosol within 15 min in control cells (Fig 6E-H). In contrast, Vav inhibition delayed A1 and A2 recruitment with limited recruitment to the membrane even after 30 min (Fig 6E-H). Based on these data, we conclude that Vav signaling is needed to recruit A1 and A2 during membrane repair.

## DISCUSSION

Here, we found that Vav2 serves as an upstream activator of membrane repair in response to cholesterol-dependent cytolysins in multiple mammalian cell types. Vav2 acted in multiple Ca^2+^-dependent repair pathways, accounting for ∼90% of Ca^2+^-dependent repair. Consistent with this large contribution to membrane repair, Vav2 coordinated microvesicle shedding via the MLK3-MEK pathway, patch repair via dysferlin, and clogging via annexin recruitment. However, we found that ceramide-mediated protection was a parallel pathway to Vav. This suggests that Vav activation is an early event during membrane repair, because it serves as a master regulator in response to bacterial CDCs.

We identified the Rac GEF that activates multiple, distinct membrane repair pathways. Our findings provide new context for prior work showing that Rac1 is activated by the CDCs pneumolysin and listeriolysin O, or laser wounding, and remodels actin upon activation (Iliev *et al*., 2007; Lam *et al*., 2018; Togo & Steinhardt, 2004). While cdc42 and Rho activation are needed for repair from laser wounding in *Xenopus* oocytes and *Drosophila* embryos (Abreu-Blanco *et al*, 2014; Benink & Bement, 2005; Moe *et al*., 2021), we did not observe that for SLO or PFO. This is consistent with prior findings that listeriolysin O damage triggered Rac (Lam *et al*., 2018). Since actin remodeling is a well-established outcome of Rac signaling, we did not focus on actin remodeling in this study. However, our findings suggest that Rac signaling has the capacity to mediate repair via multiple mechanisms beyond actin remodeling.

Distinct from prior findings focused on actin remodeling during repair, we now connect Rac activation by Vav2 to other established downstream membrane repair pathways, including MEK-dependent repair, annexins, and dysferlin activation. Future work is needed to determine if Rac1 and Rac3 are redundant, and how Rac activates these repair pathways. Since Rac binds directly to the crib domain of MLK3 (Burbelo *et al*, 1995), it could regulate MEK-dependent repair by directly activating MLK3. The mechanism by which Rac promotes dysferlin or A1 recruitment may be less direct, or Vav2 may activate them via Rac-independent mechanisms (Bustelo, 2014). Thus, our work places Rac in a central role in directing the membrane repair response to bacterial CDCs beyond its involvement in the actin cytoskeleton.

The role of other Rho GTPases remains less clear. Prior work found that Rho mediated Ca^2+^-independent repair (Iliev *et al*., 2007), and can activate non-muscle myosin II (Togo & Steinhardt, 2004). In contrast, we found that Rho sensitized cells only to SLO. Future work is needed to determine if the differences between SLO and PFO are due to membrane binding, and how this pathway integrates into the larger picture of repair. Based on our results, Rho does not appear to orchestrate a universal Ca^2+^-dependent repair mechanism.

Our work had some limitations. While our data do not support a role for cdc42 in membrane repair or for Rho in repair responding to SLO, we did not rule out activation of these proteins. These proteins could be activated, but not necessary for cell survival. We extensively used inhibitors because Rho GTPase inhibitors are well-characterized and carry the greatest translational capacity, but we did not use siRNA for Rho GTPases or GEFs. Our CRISPR cells targeted two of the three Vav2 alleles, leaving residual Vav2 expression that protected cells. Our analysis focused on SLO and PFO as representative CDCs instead of other CDCs. There is variability in toxin assays, which can complicate interpretation. We did not use a bacterial model for damage to avoid complications from non-toxin factors. Despite these limitations, we provide evidence supporting a sweeping role for Vav activation of Rac in membrane repair to bacterial CDCs.

This work opened new horizons for future study. Identification of the upstream activator for Vav2 and how it connects to Ca^2+^-influx remains to be determined. The mechanisms by which Vav2 regulates A1 and dysferlin remain to be determined. If Vav1 is redundant with Vav2, and if Vav3 antagonizes repair, remains to be determined. Another new avenue of research is the extent to which other pathways we were unable to test, like ESCRT-mediated shedding or other annexins, are controlled by Vav2. While Vav activation coordinates repair against multiple bacterial toxins, Vav may also coordinate resistance to mammalian PFTs and other toxin families. Overall, our results provide an integrated mechanism of membrane repair and suggest that triggering Vav-activated repair pathways could provide a new approach to treating NSTIs.

## MATERIALS AND METHODS

### Reagents

Unless otherwise noted, all reagents were from ThermoFisher Scientific (Waltham, MA, USA). Propidium iodide (PI) (Cat# P4170-100MG) and DAPI (Cat# D9542) were from Sigma-Aldrich (St. Louis, MO, USA). MEK inhibitor U0126 was from Tocris (Minneapolis, MN, USA) (Cat# 1144) or Cell Signaling Technology (Danvers, MA, USA) (Cat# 9903S). MLK inhibitor URMC-099 (Cat# 19147) and DL-threo-PPMP (hydrochloride) (Cat#17236) were from Cayman Chemicals (Ann Arbor, MI, USA). MEK activator PAF16 (Cat# 2940), Rac inhibitors EHT 1864 (Cat# 3872), and NSC23766 (Cat# 2161), and the Src/Syk inhibitor MNS (Cat# 2877) were from Tocris Bioscience. Rho Inhibitor-Y16 (Cat# HY-12649) and ROCK1/2 inhibitor, Y-27632 dihydrochloride (Cat# HY-10583), were from MedChemExpress (NJ, USA). Rac inhibitor EHop016 (Cat# S7319), cdc42 inhibitor ML141(Cat# S7686), and FAK/Pyk2b inhibitor PF431396 (Cat# S7644) were from Selleckchem (Houston, TX, USA). Negative control siRNA (Cat # 462001), Silencer pre-designed siRNAs for annexin A1 (Assay ID: 146988) and A2 (Assay ID: 147285) were from Ambion. PYK2/PTK2B DsiRNA were from Integrated DNA Technologies (IDT) (Coralville, IA, USA). Anti-MEK (9122L), anti–phospho–MEK-[Ser217/Ser221] (9121S), anti-ERK p44/42 (9102S), and anti–phospho–ERK [Thr202/Tyr404] (9101S) antibodies were from Cell Signaling Technologies (Danvers, MA, USA). Anti-Vav2 (Cat# 21924-1AP) was from Proteintech (Rosemont, IL). Anti-β-actin AC-15 mAb (Cat# GTX26276) was from GeneTex (Irvine, CA, USA). Anti-tubulin E7 mAb was from the Developmental Studies Hybridoma Bank, created by the NICHD of the NIH and maintained at the University of Iowa, Department of Biology (Iowa City, IA, USA). Goat anti-mouse (711-035-151) and anti-rabbit (711-035-152) horseradish peroxidase (HRP)–conjugated antibodies were from Jackson ImmunoResearch (West Grove, PA, USA).

### Plasmids

The pBAD-gIII plasmid encoding His-tagged SLO lacking the lone Cys (C530A) codon-optimized for *Escherichia coli* expression was previously described (Ray *et al*., 2022). Cys-less, His-tagged PFO in pET22 (Shepard *et al*, 1998) was a gift from Rodney Tweten (University of Oklahoma Health Sciences Center, Oklahoma City, OK, USA). Pig annexin A1 fused to YFP was a gift from Annette Draeger (University of Bern, Bern, Switzerland) (Babiychuk *et al*, 2009b). Annexin A2-GFP was a gift from Volker Gerke & Ursula Rescher (Addgene plasmid # 107196) (Rescher *et al*, 2000). OlyA fused to mCherry in pET21c(+) was a gift from Kristina. Sepčić (University of Ljubljana, Ljubljana, Slovenia) (Skocaj *et al*, 2014). The mCherry-OlyA E69A, which removes the cholesterol requirement for OlyA binding to sphingomyelin, was previously generated (Ray *et al*., 2022).

### Mice

All experimental mice were housed and maintained in accordance with Texas Tech University Institutional Animal Care and Use Committee (TTU IACUC) standards, adhering to the Guide for the Care and Use of Laboratory Animals (8th edition, NRC 2011). TTU IACUC approved mouse use. C57BL/6 mice were purchased from the Jackson Laboratory (Bar Harbor, ME, USA) (stock #000664). BMDMs were prepared using mice of both sexes aged 6 to 15 weeks as described below. No sex effects were observed. Mice were euthanized by asphyxiation through the controlled flow of pure CO_2_, followed by cervical dislocation.

### Recombinant toxins

Toxins were purified as previously described (Ray *et al*., 2022; Thapa & Keyel, 2023). Briefly, toxins were induced with 0.2% arabinose (SLO) or 0.2 mM isopropyl-β-d-thiogalactopyranoside (PFO, aerolysin) for 3 hours at room temperature, followed by purification using Nickel–NTA resin. Protein concentration was measured by Bradford assay, and hemolytic activity was determined as previously described (Romero *et al*., 2017) using human red blood cells (Zen-Bio, Research Triangle Park, NC, USA). One hemolytic unit (HU) is the amount of toxin needed to lyse 50% of a 2% human red blood cell solution in 30 min at 37°C in phosphate-buffered saline (PBS) with 2 mM CaCl_2_, 0.3% bovine serum albumin, and 10 mM Hepes (pH 7.4) (Keyel *et al*, 2012). Hemolytic units per milliliter (HU/ml) were used to normalize toxin activities and achieve coherent cytotoxicity across toxin preparations (Supplementary Table S1).

### Cell culture

HeLa cells (ATCC (Manassas, VA, USA) CCL-2), and HEK-293 (ATCC CRL-1573) were cultured in Dulbecco’s modified Eagle’s medium (DMEM; Corning, Corning, NY, USA) supplemented with 10% Equafetal bovine serum (Atlas Biologicals, Fort Collins, CO, USA), 1× l-glutamine, 1×penicillin and streptomycin (D10). B6 bone marrow–derived macrophages (BMDMs) were isolated from bone marrow and cultured as previously described (Ray *et al*., 2022; Romero *et al*., 2017). Briefly, BMDMs were differentiated for 7–21 days in DMEM supplemented with 30% L929 cell supernatant, 20% fetal calf serum (VWR Seradigm, Radnor, PA, USA), 1 mM sodium pyruvate, and 1× L-glutamine. C2C12 myoblasts stably expressing control (ATCC CRL-3419) or Dysf-shRNA (ATCC CRL-3418) were cultured in D10 plus 2 μg/ml puromycin. Cells were maintained at 37°C in 5% CO_2_.

### Transfection

For siRNA knockdowns, HeLa cells were plated at a density of 2 × 10^5^ cells in six-well plates. After 24 hours, these cells were transfected with 20 nM control, Pyk2, annexins A1, or A2 siRNAs using Lipofectamine 2000 in Opti-MEM for 48 to 72 hours prior to assays.

For plasmid transfections, HeLa cells were plated at 2 × 10^5^ cells per 35-mm glass-bottom dish, 35-mm dish, or per well of a six-well plate. They were then transfected with 500 ng of annexin A1-YFP or 750 ng of annexin A2-GFP using Lipofectamine 2000 in Opti-MEM 2 days before imaging. The D10 was replaced on the day following transfection. Transfection efficiency for each construct ranged from 50-70% across experiments.

### CRISPR knockout of Vav2

HeLa cells were plated at 2.5 × 10^5^ cells in six-well plates. After 24 hours, these cells were transfected with either negative control guide RNA or guide RNA targeting Vav2, along with 2.5 μg TrueCut Cas9 (Cat A36497) using CrisprMax in Opti-MEM. After 4 days, cells were harvested and plated at 0.6 cells/well in 96-well plates for single cell cloning. Remaining cells were assayed for CRISPR efficiency using the GeneArt Genomic Cleavage Detection kit per manufacturer’s instructions. Wells with single colonies were expanded and screened by western blot to determine the extent of Vav2 expression. Control clones and clones with reduced Vav2 expression were then assayed for cytotoxicity, Vav2 expression, and cryopreserved.

### Cytotoxicity assays

Cytotoxicity assays were performed as previously described (Keyel *et al*., 2011; Ray *et al*., 2022; Ray *et al*, 2018; Thapa & Keyel, 2023), with the following modifications. 1 ×10^6^ HeLa cells were plated in 60 mm dishes and incubated at 37°C and 5% CO_2_ incubator for 36-48 hours before the cytotoxicity assay. At 80-95% confluency, media was aspirated, cells were washed with 1X PBS and serum-starved in DMEM for 30 min at 37°C. Cells were then treated with 20 μM inhibitors unless otherwise noted before toxin challenge. DMSO was used as the vehicle control, ML141 inhibits cdc42, Y16 inhibits Rho, Y27632 inhibits ROCK1/2, URMC099 inhibits MLK3, U0126 inhibits MEK, EGTA chelates Ca^2+^, EHop016, EHT1864, and NSC23766 inhibit Rac, PF431396 inhibits FAK and Pyk2b, and MNS inhibits Src and Syk. Cells were incubated with inhibitors in DMEM at 37°C, 5% CO_2_ for 30 min before toxin challenge.

Then cells were harvested, and 1 × 10^5^ pretreated cells were challenged in suspension with 31-2000 HU/mL toxin for 30 min at 37°C in RPMI supplemented with 2 mM CaCl_2_ (RC) and 20 μg/ml propidium iodide (PI) in the continued presence of inhibitors or DMSO. Cells were analyzed on a four-laser Attune Nxt flow cytometer. Debris was gated out, and the percentage of single cells with high dye (PI) fluorescence (2 to 3 log shift) (dye high) was quantified. We previously showed that using this gating and selecting “dye high” populations accurately reports dead cells (Keyel *et al*., 2011; Ray *et al*., 2018). We calculated % specific lysis as follows:

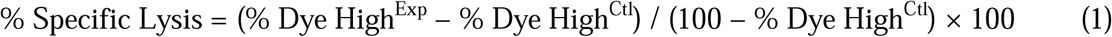

The toxin dose required to kill 50% of cells was defined as the LC_50_. The LC_50_ was determined by logistic modelling using Excel (Microsoft, Redmond, WA, USA) as previously described (Haram *et al*., 2022). The logistic model used was:

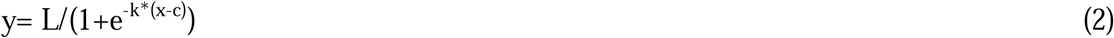

where x is the toxin concentration, y is the % specific lysis, L is maximal specific lysis, c is the toxin concentration at which y is 50% of L, and k measures the steepness of the sigmoidal curve.

### SDS-PAGE and Immunoblotting

We performed SDS-PAGE as previously described (Ray *et al*., 2022; Thapa & Keyel, 2023). Briefly, cells were spun down and washed thrice with 1X PBS before resuspension in 95°C 1X SDS sample buffer, incubation at 95°C for 5 min and sonication. Samples were resolved on 10% polyacrylamide gels and transferred to nitrocellulose in an ice bath with transfer buffer at 110 V for 90 min. Blots were blocked using 5% skim milk in Tris-buffered saline with tween (TBST) for 2 hours. Portions of the blots were incubated for 2 h at room temperature with any of the following primary antibodies diluted in 1% skim milk in TBST: CPTC-A1–3 anti-annexin A1 (1:250) mAb, anti-Annexin A2 (1:1000) mAb, AC-15 anti-β-actin (1:5000) mAb, rabbit polyclonal antibodies: anti-MEK, anti–phospho-MEK [Ser217/221], anti-ERK p44/42, anti–phospho-ERK [Thr202/Tyr404] at 1:1000 dilution each, and anti-PYK2 (1:250). After washing, blots were incubated with HRP-conjugated anti-mouse or anti-rabbit immunoglobulin G antibodies (1:10,000) diluted in 1% skim milk and developed with enhanced chemiluminescence (ECL): 0.01% H_2_O_2_ (Walmart, Fayetteville, AR, USA), 0.2 mM p-Coumaric acid (Sigma-Aldrich), 1.25 mM luminol (Sigma-Aldrich) in 0.1 M Tris (pH 8.4). Blots from at least three independent experiments were quantified by measuring and comparing band intensities using Photoshop (Adobe, San Jose, CA, USA). Complete western blots are included in the supplemental data (Fig S6).

### Immunofluorescence

Immunofluorescence was performed as previously described (Thapa & Keyel, 2023). HeLa cells were plated on coverslips. Cells were washed twice in 1X PBS, fixed in 2% paraformaldehyde (PFA) for 15 min, washed, permeabilized and blocked with 0.2% saponin, 10% goat serum in PBS, stained with phalloidin conjugated to Alexa Fluor 488 (1:40) for one hour, washed, DAPI stained, washed, and mounted on slides in gelvatol. For Vav2 expression, anti-Vav2 (1:200) and anti-tubulin (1:10) were used, with anti-mouse IgG conjugated to Alexa Fluor 488 (1:500) and anti-rabbit IgG conjugated to Alexa Fluor 568 (1:500) as secondary antibodies. Transfected cells were fixed and DAPI stained as described above. Cells were imaged on a Fluoview 3000 confocal microscope (Olympus, Tokyo, Japan) with a 60×, 1.42 NA oil-immersion objective.

Images were processed using ImageJ (NIH, Bethesda, MD, USA).

### Statistics

GraphPad Prism 10.0.3 (San Diego, CA, USA) was used for statistical analysis. Statistical significance was determined by one-way analysis of variance (ANOVA) or repeated-measures ANOVA with Tukey post hoc testing. p< 0.05 was statistically significant.

## Supporting information

Full Blots

## ACKNOWLEDGMENTS

The authors thank the Keyel lab members for their critical review of the manuscript and the College of Arts & Sciences Microscopy for using their facilities. This work was supported by the National Institute of Allergy and Infectious Diseases of the National Institutes of Health grant R21AI156225 (PAK) and American Heart Association Innovative Project Award 23IPA1053083 (https://doi.org/10.58275/AHA.23IPA1053083.pc.gr.171946) (PAK).

## AUTHOR CONTRIBUTIONS

VGK: methodology development, investigation, data curation, formal analysis, visualization, writing—original draft, writing—review & editing.

AH: investigation, data curation, writing—original draft, writing—review & editing.

TK: investigation, data curation, writing—review & editing.

PAK: Conceptualization, methodology development, writing—original draft, data curation, formal analysis, writing—review & editing, supervision, project administration, funding acquisition.

## ETHICAL APPROVAL

All mouse experimental procedures were approved and in accordance with the Texas Tech University Institutional Animal Care and Use Committee (TTU IACUC).

## AVAILABILITY OF DATA AND MATERIALS

All data are in the main text or supplementary materials.

**Graphic Abstract. Vav is a master regulator of membrane repair in response to bacterial pore-forming toxins.** Upon pore formation and membrane damage by cholesterol-dependent cytolysins (CDCs) like streptolysin O and perfringolysin O, several membrane repair pathways are activated due to calcium ion (Ca^2+^) influx. Ca^2+^ influx activates the Rac GEF Vav2, which can activate Rac. Downstream of this activation, at least three pathways are activated: 1) Membrane shedding is activated by the MLK3-MEK-Annexin A2 pathway. 2) Patch repair by dysferlin, and 3) clogging by other annexins like Annexin A1. In contrast, ceramide-mediated protection is parallel to the Vav2 pathways.

## Supplementary Tables and Figures

**Supplementary Table S1.** Specific activity and protein concentration of toxins used.

| Toxin | Figure used | Hemolytic activity (HU/mL) | Protein conc (mg/mL) | Specific activity (HU/mg) |
| --- | --- | --- | --- | --- |
| SLO WT | 1A-D, 2A-D, S1B-H, S2A-D | $1.2 \times 10^6$ | 2.7 | $4.4 \times 10^6$ |
| SLO WT | 3A, 4A-B, 4D, 5A-F, S3B-E, S4, S5A | $2.0 \times 10^6$ | 5.3 | $3.8 \times 10^5$ |
| SLO WT | 1E-F, 6B, 6D-F, S5A-B | $4.8 \times 10^6$ | 2.7 | $1.8 \times 10^6$ |
| PFO WT | 1A-F, 2A-D, 3A, 4A-B, 4D, 5A-C, 5F, 5H, 6B, 6D, S1-4A-D | $2.0 \times 10^6$ | 5.6 | $3.6 \times 10^5$ |
| PFO WT | 1E-G, S4E, S5A-B | $6.4 \times 10^6$ | 7.3 | $8.8 \times 10^5$ |
| Aerolysin WT | S1H | $4.0 \times 10^5$ | 2 | $2.0 \times 10^5$ |

**Supplementary Fig S1. Rac inhibition sensitizes cells to bacterial pore-forming toxins.** (A) HeLa cells plated on coverslips were serum-starved for 30 min, treated with DMSO, or 20 μM of the indicated inhibitor for 30 min, fixed, stained with phalloidin Alexa flour 488 (green) and DAPI (blue), and imaged by confocal microscopy. Arrowheads point to actin cytoskeleton or its disruption. Micrographs represent three independent experiments. Scale bar=10 μm. (B) Actin content from the micrographs was quantitated to show total and mean F-actin content. (C-I) HeLa cells were pretreated with vehicle (DMSO) or 5, 10, 20 μM (C) EHop016, (D) ML141, (E) Y16, (F) Y27632, or (G-I) 20 μM ML141, Y16, Y27632, and/or EHop016 for 30 min. Cells were then challenged with the indicated concentrations of (C-H) SLO, PFO, or (I) aerolysin for 30 min at 37°C. PI uptake was analyzed by flow cytometry. The LC_50_ was calculated as described in the methods. The dotted line indicates the limit of detection. Points on this line had LC_50_>1000 HU/mL. Graphs show (I) individual experiments with mean ±SEM, or (C-H) the mean ±SEM for n=5 (C, D, E, (SLO), D, E, (PFO)), n=4 (C, G, (PFO), G, H, (SLO), I, (Aero)), n=6 (E, SLO), or n=3 (H, PFO). The dotted line indicates the limit of detection. Points on this line had LC_50_>1000 HU/mL (I). *p<0.05, **** p<0.0001 denotes statistical significance using repeated-measures ANOVA between groups with Tukey post-test.

**Supplementary Fig S2. Multiple cell types are sensitive to Rac inhibition.** (A) HeLa, (B) bone marrow-derived macrophages (B6 M), (C) HEK, or (D) C2C12 cells were serum-starved for 30 min, treated with DMSO or 5, 10, or 20 μM (A) Rac inhibitor EHT1864, (B) 20 μM EHT1864, or (C, D) 20 μM EHop016 for 30 min, challenged with the indicated concentrations of SLO or PFO, and analyzed by flow cytometry. Graphs display the mean ±S.E.M for n=3 (A-D) or n=4 (A, PFO) independent experiments.

**Supplementary Fig S3. Vav2 promotes membrane repair independent of the tyrosine kinases Src, Syk, focal adhesion kinase, and Pyk2.** (A) HeLa cells plated on coverslips were serum-starved for 30 min, treated with DMSO or 100 μM NSC23766 for 30 min, fixed, stained with phalloidin Alexa Fluor 488 (green) and DAPI (blue), and imaged by confocal microscopy. Arrowheads point to actin cytoskeleton or its disruption. (B) HeLa cells were serum-starved for 30 min, treated with DMSO or 50, 100 μM NSC23766 and challenged with the indicated concentrations of SLO or PFO for 30 min at 37°C. Propidium iodide (PI) uptake was analyzed by flow cytometry. (C) Untransfected HeLa cells, CRISPR Control clone (CTL), or Vav2 CRISPR clones (18, 36, and 44) were challenged with the indicated concentrations of SLO for 30 min at 37°C and analyzed as in (B). (D, E) HeLa cells were treated as in (B) using (D) 5, 10, or 20 μM Src/Syk inhibitor MNS or (E) 5, 10, 20 μM FAK/Pyk2 inhibitor PF431396. (F) Control or Pyk2B-siRNA transfected HeLa cells were treated with DMSO or 20 μM EHop016 for 30 min and then challenged with the indicated concentrations of SLO or PFO for 30 min. Graphs show the mean ±S.E.M for n=4 (B, D, SLO) or n=3 ((C, E, F, (SLO)), (B, D, E, F (PFO))). Micrographs represent three independent experiments. Scale bar=10 μm.

**Supplementary Fig S4. Vav2 promotes Ca^2+^-dependent repair upstream of MLK3-MEK signaling but parallel to ceramide-mediated repair.** (A-D) HeLa cells were serum-starved for 30 min and treated with DMSO, 20 μM EHop016, (B-C) U0126, (C) URMC099, and/or (D) 20 μM PAF16 for 30 min and then challenged with the indicated concentrations of SLO or PFO for 30 min. Where indicated, 2 mM EGTA was used instead of 2 mM CaCl_2_. (E) HeLa cells were treated with 20 μM PPMP for 72 h at 37°C, then treated with DMSO or 20 μM EHop016, challenged and analyzed as in (A).Graphs display the mean ±S.E.M for n=3 (A, B, D, (SLO), A, B, C, D, E, (PFO)) or n=4 (C, SLO) or n=5 (E, SLO).

**Supplementary Fig S5. Vav regulates dysferlin and annexin-mediated repair.** (A) Dysferlin-shRNA or control shRNA C2C12 or (B) HeLa cells transfected with control (Ctrl), annexin A1 (A1), or annexin A2 (A2) siRNA were serum-starved for 30 min, treated with DMSO or 20 μM EHop016 for 30 min and then challenged with the indicated concentrations of SLO or PFO for 30 min. PI uptake was analyzed by flow cytometry. Graphs show the mean ±SEM for n=4 (A (SLO)), or n=3 (B, (SLO) A, B, (PFO)).

## REFERENCES

1. Abreu-Blanco MT, Verboon JM, Parkhurst SM (2014) Coordination of Rho family GTPase activities to orchestrate cytoskeleton responses during cell wound repair. Curr Biol 24: 144–155

2. Alves S, Pereira JM, Mayer RL, Goncalves ADA, Impens F, Cabanes D, Sousa S (2022) Cells Responding to Closely Related Cholesterol-Dependent Cytolysins Release Extracellular Vesicles with a Common Proteomic Content Including Membrane Repair Proteins. Toxins (Basel) 15

3. Awad MM, Ellemor DM, Boyd RL, Emmins JJ, Rood JI (2001) Synergistic effects of alpha-toxin and perfringolysin O in Clostridium perfringens-mediated gas gangrene. Infect Immun 69: 7904–7910

4. Babiychuk EB, Monastyrskaya K, Potez S, Draeger A (2009a) Intracellular Ca(2+) operates a switch between repair and lysis of streptolysin O-perforated cells. Cell Death Differ 16: 1126–1134

5. Babiychuk EB, Monastyrskaya K, Potez S, Draeger A (2009b) Intracellular Ca(2+) operates a switch between repair and lysis of streptolysin O-perforated cells. Cell Death Differ 16: 1126–1134

6. Babiychuk EB, Monastyrskaya K, Potez S, Draeger A (2011) Blebbing confers resistance against cell lysis. Cell Death Differ 18: 80–89

7. Benink HA, Bement WM (2005) Concentric zones of active RhoA and Cdc42 around single cell wounds. J Cell Biol 168: 429–439

8. Boye TL, Maeda K, Pezeshkian W, Sonder SL, Haeger SC, Gerke V, Simonsen AC, Nylandsted J (2017) Annexin A4 and A6 induce membrane curvature and constriction during cell membrane repair. Nat Commun 8: 1623

9. Burbelo PD, Drechsel D, Hall A (1995) A conserved binding motif defines numerous candidate target proteins for both Cdc42 and Rac GTPases. J Biol Chem 270: 29071–29074

10. Bustelo XR (2014) Vav family exchange factors: an integrated regulatory and functional view. Small GTPases 5: 9

11. Chakrabarti S, Kobayashi KS, Flavell RA, Marks CB, Miyake K, Liston DR, Fowler KT, Gorelick FS, Andrews NW (2003) Impaired membrane resealing and autoimmune myositis in synaptotagmin VII-deficient mice. J Cell Biol 162: 543–549

12. Cooper ST, McNeil PL (2015) Membrane Repair: Mechanisms and Pathophysiology. Physiol Rev 95: 1205–1240

13. Dany M, Ogretmen B (2015) Ceramide induced mitophagy and tumor suppression. Biochim Biophys Acta 1853: 2834–2845

14. Demonbreun AR, Allen MV, Warner JL, Barefield DY, Krishnan S, Swanson KE, Earley JU, McNally EM (2016a) Enhanced Muscular Dystrophy from Loss of Dysferlin Is Accompanied by Impaired Annexin A6 Translocation after Sarcolemmal Disruption. Am J Pathol 186: 1610–1622

15. Demonbreun AR, Quattrocelli M, Barefield DY, Allen MV, Swanson KE, McNally EM (2016b) An actin-dependent annexin complex mediates plasma membrane repair in muscle. J Cell Biol 213: 705–718

16. Du Y, Bock BC, Schachter KA, Chao M, Gallo KA (2005) Cdc42 induces activation loop phosphorylation and membrane targeting of mixed lineage kinase 3. J Biol Chem 280: 42984–42993

17. Gao Y, Dickerson JB, Guo F, Zheng J, Zheng Y (2004) Rational design and characterization of a Rac GTPase-specific small molecule inhibitor. Proc Natl Acad Sci U S A 101: 7618–7623

18. Hakkarainen TW, Kopari NM, Pham TN, Evans HL (2014) Necrotizing soft tissue infections: review and current concepts in treatment, systems of care, and outcomes. Current problems in surgery 51: 344–362

19. Han S, Mistry A, Chang JS, Cunningham D, Griffor M, Bonnette PC, Wang H, Chrunyk BA, Aspnes GE, Walker DP et al (2009) Structural characterization of proline-rich tyrosine kinase 2 (PYK2) reveals a unique (DFG-out) conformation and enables inhibitor design. J Biol Chem 284: 13193–13201

20. Haram CS, Moitra S, Keane R, Breslav E, Zhang K, Keyel PA (2022) Deciphering the Molecular Mechanism and Function of Pore-Forming Toxins using Leishmania major. J Vis Exp: e64341

21. Haram CS, Moitra S, Keane R, Kuhlmann FM, Frankfater C, Hsu FF, Beverley SM, Zhang K, Keyel PA (2023) The sphingolipids ceramide and inositol phosphorylceramide protect the Leishmania major membrane from sterol-specific toxins. J Biol Chem 299: 104745

22. Iliev AI, Djannatian JR, Nau R, Mitchell TJ, Wouters FS (2007) Cholesterol-dependent actin remodeling via RhoA and Rac1 activation by the Streptococcus pneumoniae toxin pneumolysin. Proc Natl Acad Sci U S A 104: 2897–2902

23. Jabbour G, El-Menyar A, Peralta R, Shaikh N, Abdelrahman H, Mudali IN, Ellabib M, Al-Thani H (2016) Pattern and predictors of mortality in necrotizing fasciitis patients in a single tertiary hospital. World J Emerg Surg 11: 40

24. Jimenez AJ, Maiuri P, Lafaurie-Janvore J, Divoux S, Piel M, Perez F (2014) ESCRT machinery is required for plasma membrane repair. Science 343: 1247136

25. Jimenez AJ, Perez F (2017) Plasma membrane repair: the adaptable cell life-insurance. Curr Opin Cell Biol 47: 99–107

26. Kayejo VG, Fellner H, Thapa R, Keyel PA (2023) Translational implications of targeting annexin A2: From membrane repair to muscular dystrophy, cardiovascular disease and cancer. Clin Transl Discov 3

27. Keyel PA, Heid ME, Watkins SC, Salter RD (2012) Visualization of bacterial toxin induced responses using live cell fluorescence microscopy. J Vis Exp: e4227

28. Keyel PA, Loultcheva L, Roth R, Salter RD, Watkins SC, Yokoyama WM, Heuser JE (2011) Streptolysin O clearance through sequestration into blebs that bud passively from the plasma membrane. J Cell Sci 124: 2414–2423

29. Lam JGT, Vadia S, Pathak-Sharma S, McLaughlin E, Zhang X, Swanson J, Seveau S (2018) Host cell perforation by listeriolysin O (LLO) activates a Ca(2+)-dependent cPKC/Rac1/Arp2/3 signaling pathway that promotes Listeria monocytogenes internalization independently of membrane resealing. Mol Biol Cell 29: 270–284

30. Landry JJ, Pyl PT, Rausch T, Zichner T, Tekkedil MM, Stutz AM, Jauch A, Aiyar RS, Pau G, Delhomme N et al (2013) The genomic and transcriptomic landscape of a HeLa cell line. G3 (Bethesda) 3: 1213–1224

31. Limbago B, Penumalli V, Weinrick B, Scott JR (2000) Role of streptolysin O in a mouse model of invasive group A streptococcal disease. Infect Immun 68: 6384–6390

32. Los FC, Randis TM, Aroian RV, Ratner AJ (2013) Role of pore-forming toxins in bacterial infectious diseases. Microbiology and molecular biology reviews : MMBR 77: 173–207

33. Maldonado MDM, Medina JI, Velazquez L, Dharmawardhane S (2020) Targeting Rac and Cdc42 GEFs in Metastatic Cancer. Front Cell Dev Biol 8: 201

34. McNeil AK, Rescher U, Gerke V, McNeil PL (2006) Requirement for annexin A1 in plasma membrane repair. J Biol Chem 281: 35202–35207

35. McNeil PL, Kirchhausen T (2005) An emergency response team for membrane repair. Nat Rev Mol Cell Biol 6: 499–505

36. Moe A, Holmes W, Golding AE, Zola J, Swider ZT, Edelstein-Keshet L, Bement W (2021) Cross-talk-dependent cortical patterning of Rho GTPases during cell repair. Mol Biol Cell 32: 1417–1432

37. Montalvo-Ortiz BL, Castillo-Pichardo L, Hernandez E, Humphries-Bickley T, De la Mota-Peynado A, Cubano LA, Vlaar CP, Dharmawardhane S (2012) Characterization of EHop-016, novel small molecule inhibitor of Rac GTPase. J Biol Chem 287: 13228–13238

38. Nygard Skalman L, Holst MR, Larsson E, Lundmark R (2018) Plasma membrane damage caused by listeriolysin O is not repaired through endocytosis of the membrane pore. Biology open 7

39. Ray S, Roth R, Keyel PA (2022) Membrane repair triggered by cholesterol-dependent cytolysins is activated by mixed lineage kinases and MEK. Science advances 8: eabl6367

40. Ray S, Thapa R, Keyel PA (2018) Multiple Parameters Beyond Lipid Binding Affinity Drive Cytotoxicity of Cholesterol-Dependent Cytolysins. Toxins (Basel) 11

41. Rescher U, Zobiack N, Gerke V (2000) Intact Ca(2+)-binding sites are required for targeting of annexin 1 to endosomal membranes in living HeLa cells. J Cell Sci 113 (Pt 22): 3931–3938

42. Romero M, Keyel M, Shi G, Bhattacharjee P, Roth R, Heuser JE, Keyel PA (2017) Intrinsic repair protects cells from pore-forming toxins by microvesicle shedding. Cell Death Differ 24: 798–808

43. Roostalu U, Strahle U (2012) In vivo imaging of molecular interactions at damaged sarcolemma. Dev Cell 22: 515–529

44. Schoenauer R, Larpin Y, Babiychuk EB, Drucker P, Babiychuk VS, Avota E, Schneider-Schaulies S, Schumacher F, Kleuser B, Koffel R et al (2019) Down-regulation of acid sphingomyelinase and neutral sphingomyelinase-2 inversely determines the cellular resistance to plasmalemmal injury by pore-forming toxins. FASEB J 33: 275–285

45. Shepard LA, Heuck AP, Hamman BD, Rossjohn J, Parker MW, Ryan KR, Johnson AE, Tweten RK (1998) Identification of a membrane-spanning domain of the thiol-activated pore-forming toxin Clostridium perfringens perfringolysin O: an alpha-helical to beta-sheet transition identified by fluorescence spectroscopy. Biochemistry 37: 14563–14574

46. Shutes A, Onesto C, Picard V, Leblond B, Schweighoffer F, Der CJ (2007) Specificity and mechanism of action of EHT 1864, a novel small molecule inhibitor of Rac family small GTPases. J Biol Chem 282: 35666–35678

47. Skocaj M, Resnik N, Grundner M, Ota K, Rojko N, Hodnik V, Anderluh G, Sobota A, Macek P, Veranic P et al (2014) Tracking cholesterol/sphingomyelin-rich membrane domains with the ostreolysin A-mCherry protein. PLoS One 9: e92783

48. Stefl M, Takamiya M, Middel V, Tekpinar M, Nienhaus K, Beil T, Rastegar S, Strahle U, Nienhaus GU (2024) Caveolae disassemble upon membrane lesioning and foster cell survival. iScience 27: 108849

49. Swaggart KA, Demonbreun AR, Vo AH, Swanson KE, Kim EY, Fahrenbach JP, Holley-Cuthrell J, Eskin A, Chen Z, Squire K et al (2014) Annexin A6 modifies muscular dystrophy by mediating sarcolemmal repair. Proc Natl Acad Sci U S A 111: 6004–6009

50. Thapa R, Keyel PA (2023) Patch repair protects cells from the small pore-forming toxin aerolysin. J Cell Sci 136

51. Thapa R, Ray S, Keyel PA (2020) Interaction of Macrophages and Cholesterol-Dependent Cytolysins: The Impact on Immune Response and Cellular Survival. Toxins (Basel) 12

52. Togo T, Steinhardt RA (2004) Nonmuscle myosin IIA and IIB have distinct functions in the exocytosis-dependent process of cell membrane repair. Mol Biol Cell 15: 688–695

53. Wang WY, Wu YC, Wu CC (2006) Prevention of platelet glycoprotein IIb/IIIa activation by 3,4-methylenedioxy-beta-nitrostyrene, a novel tyrosine kinase inhibitor. Mol Pharmacol 70: 1380–1389

