## Supplementary figures and images for "Vav2 is a master regulator of repair against bacterial pore-forming toxins"

### Full Blots

#### Figure 3B

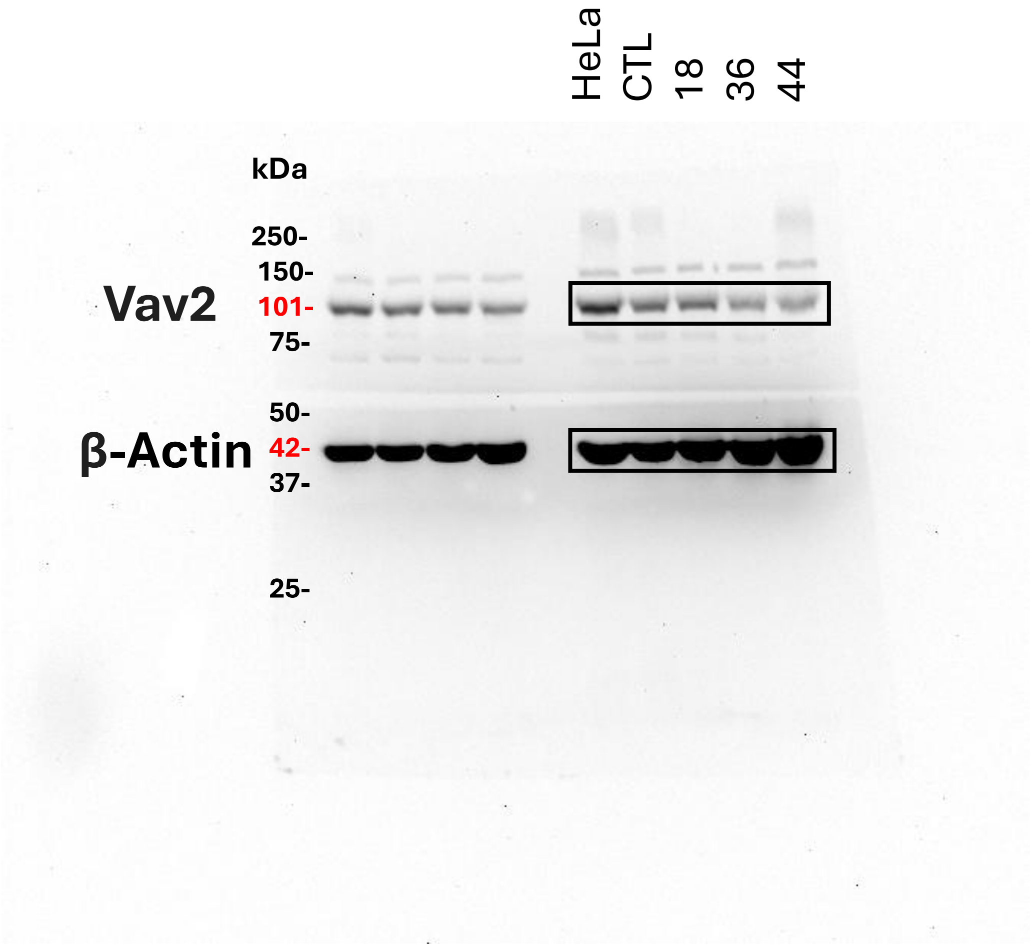

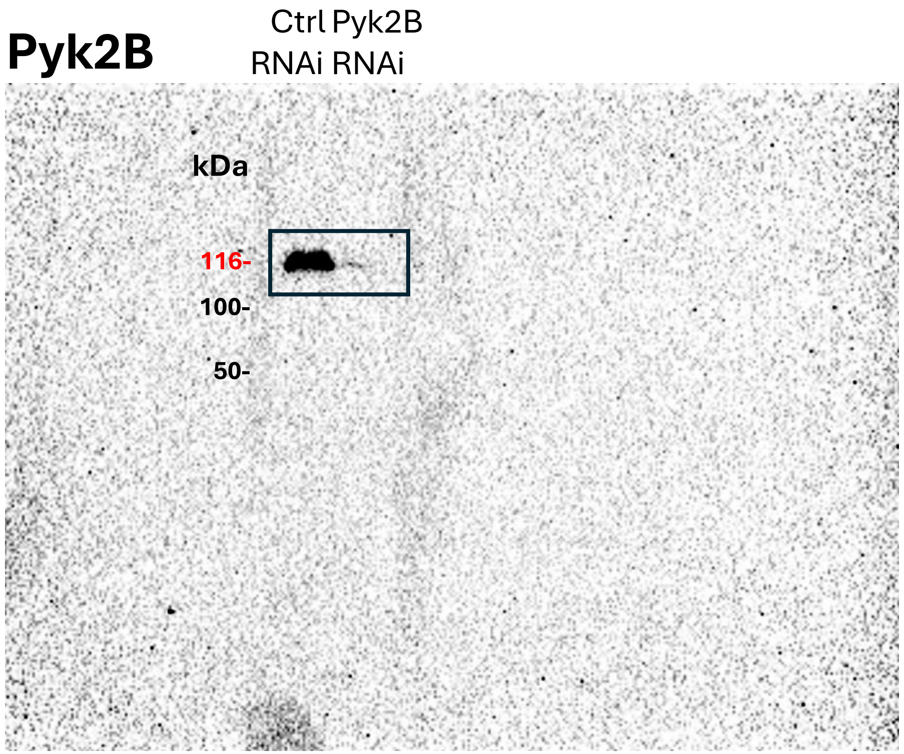

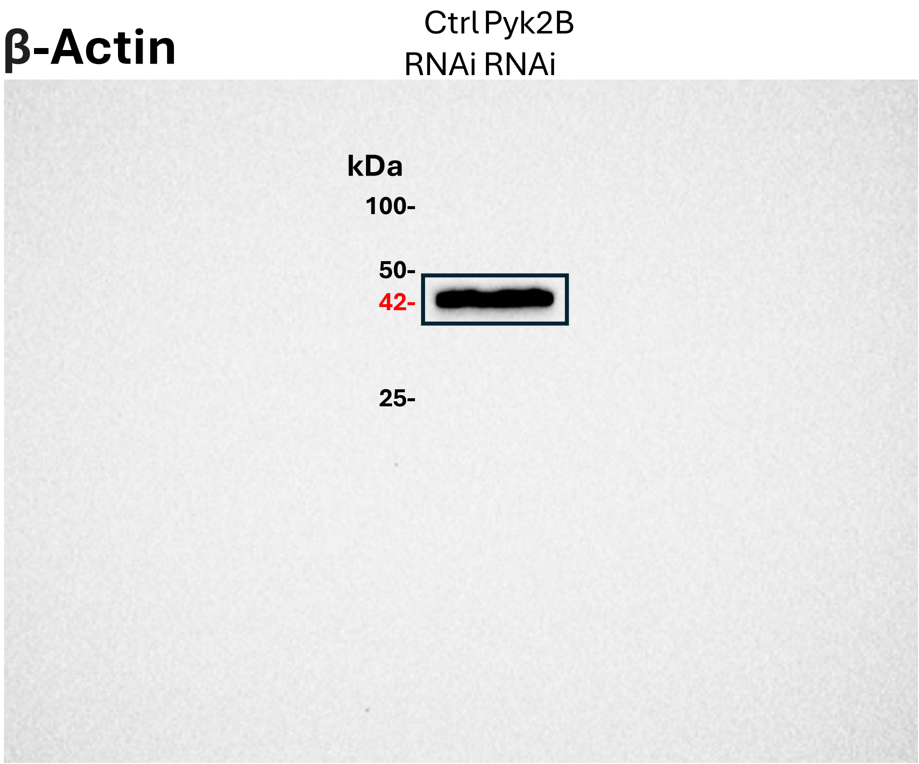

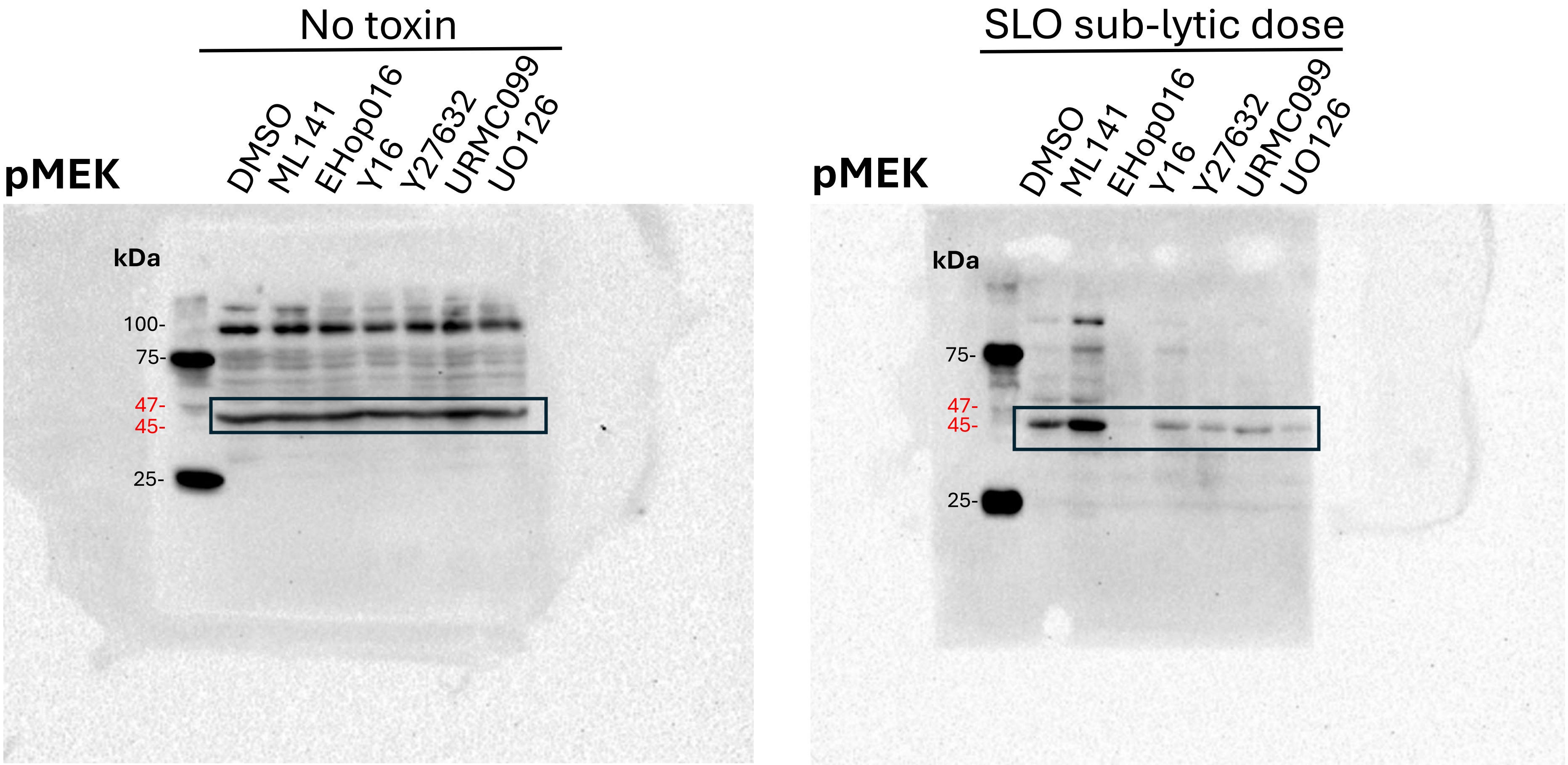

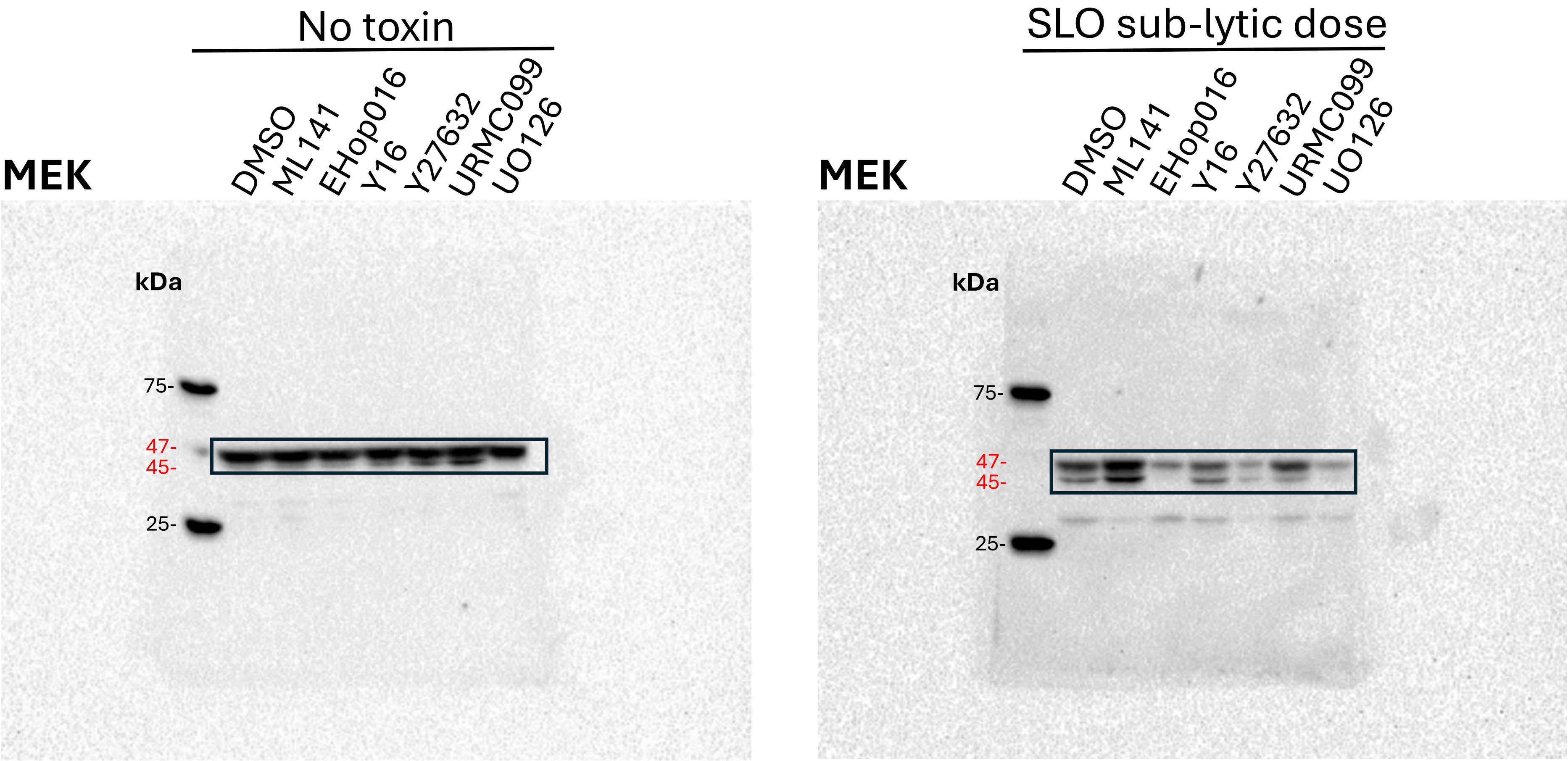

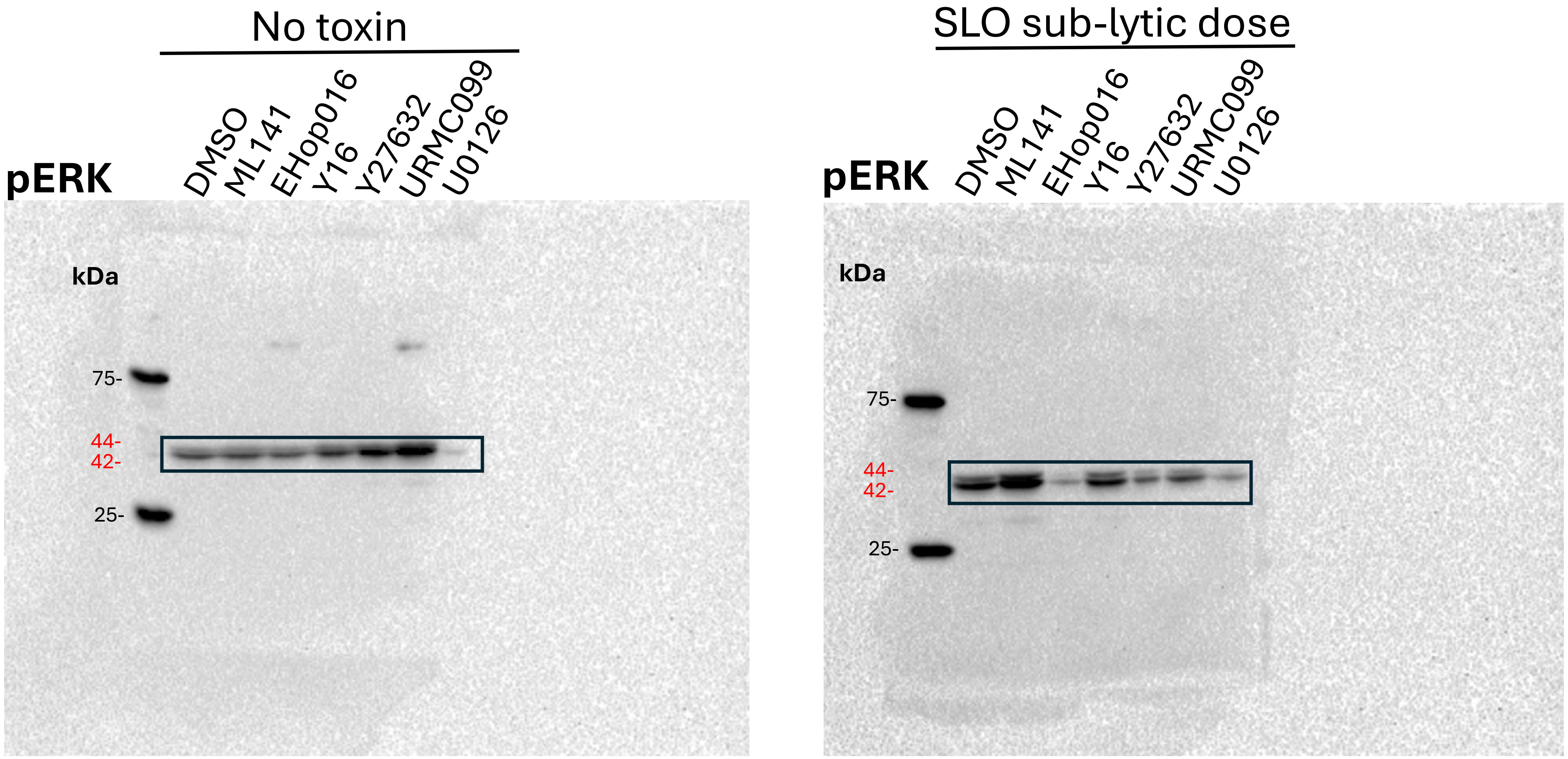

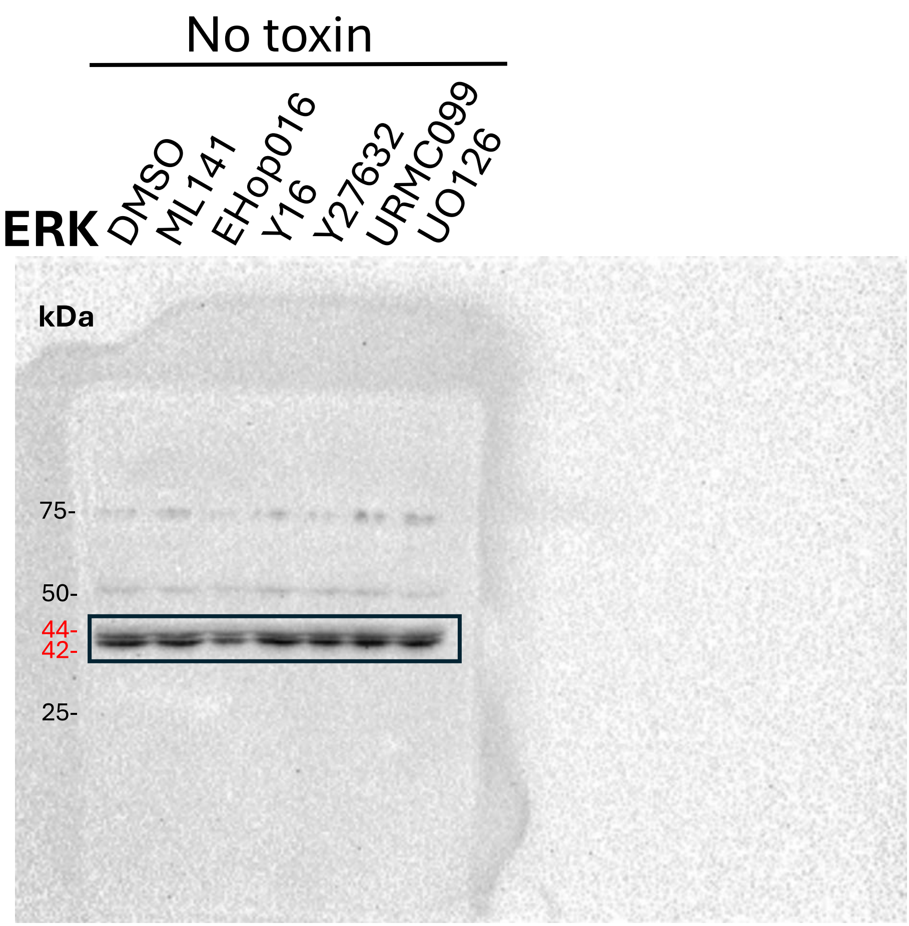

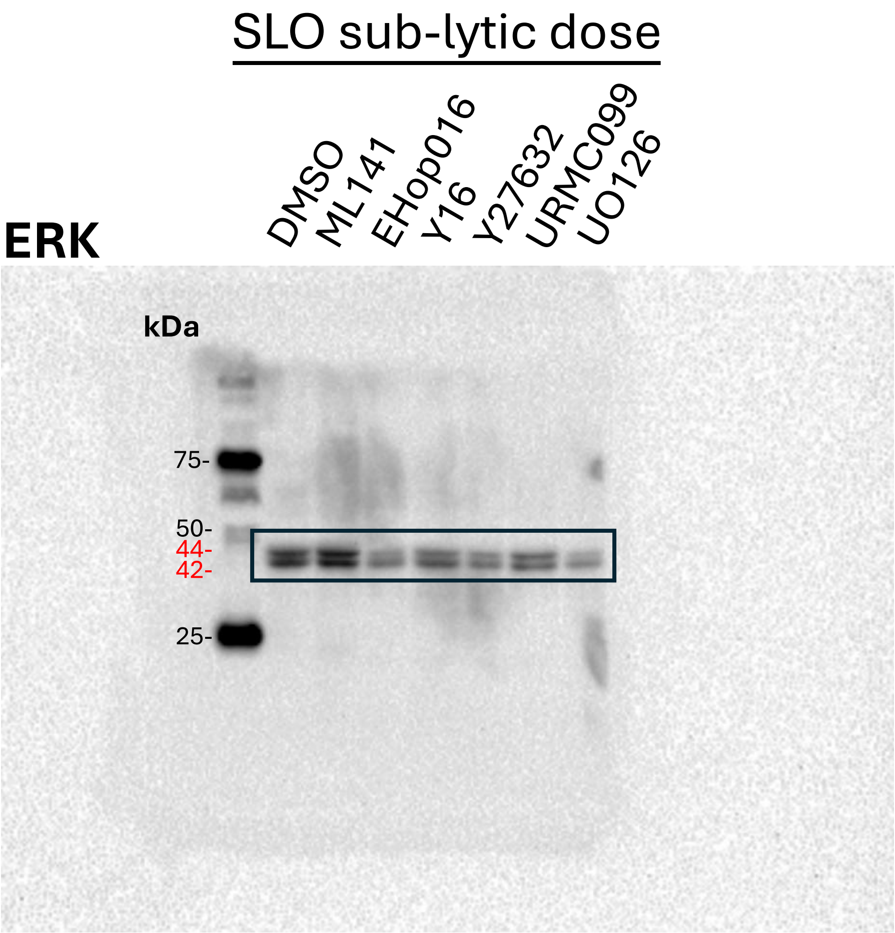

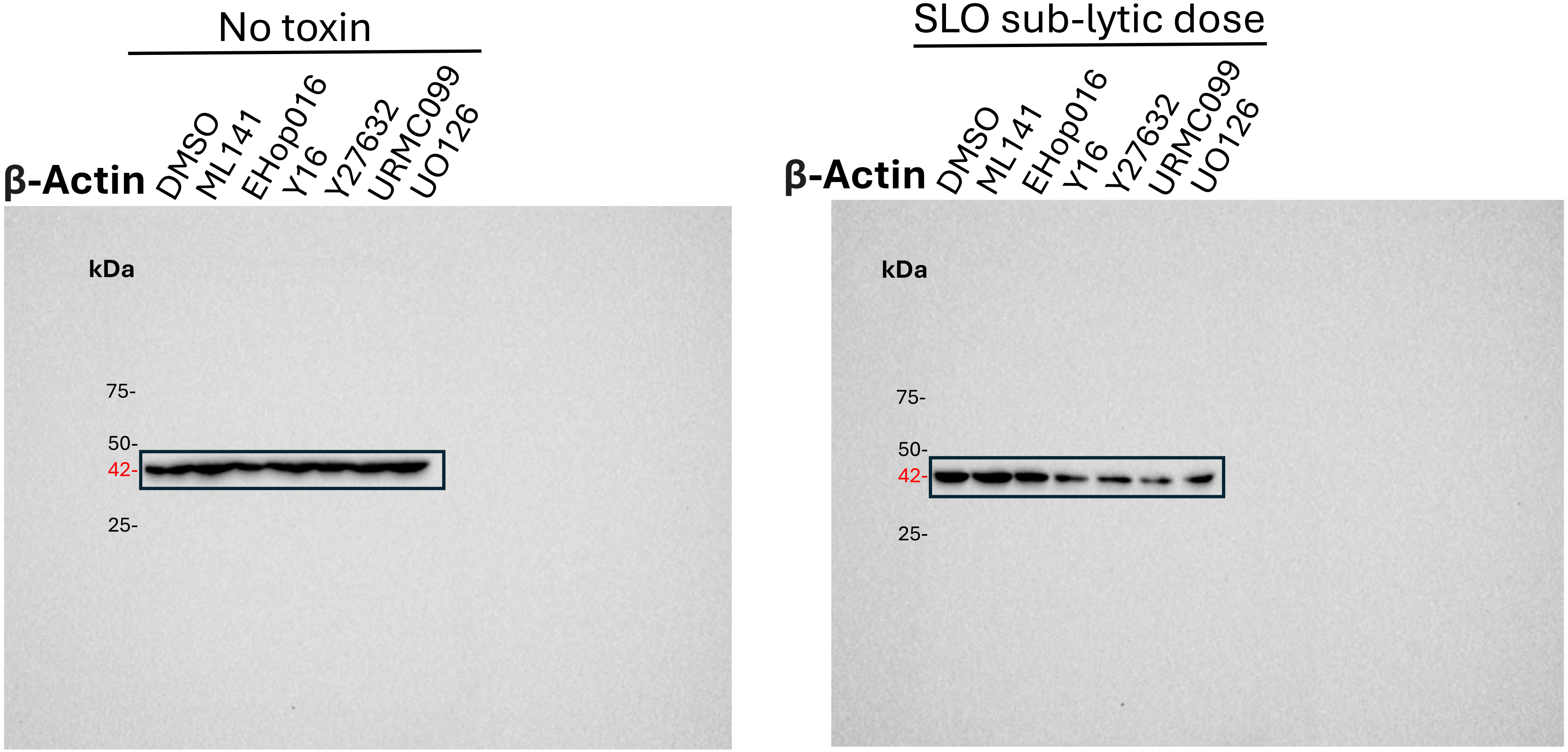

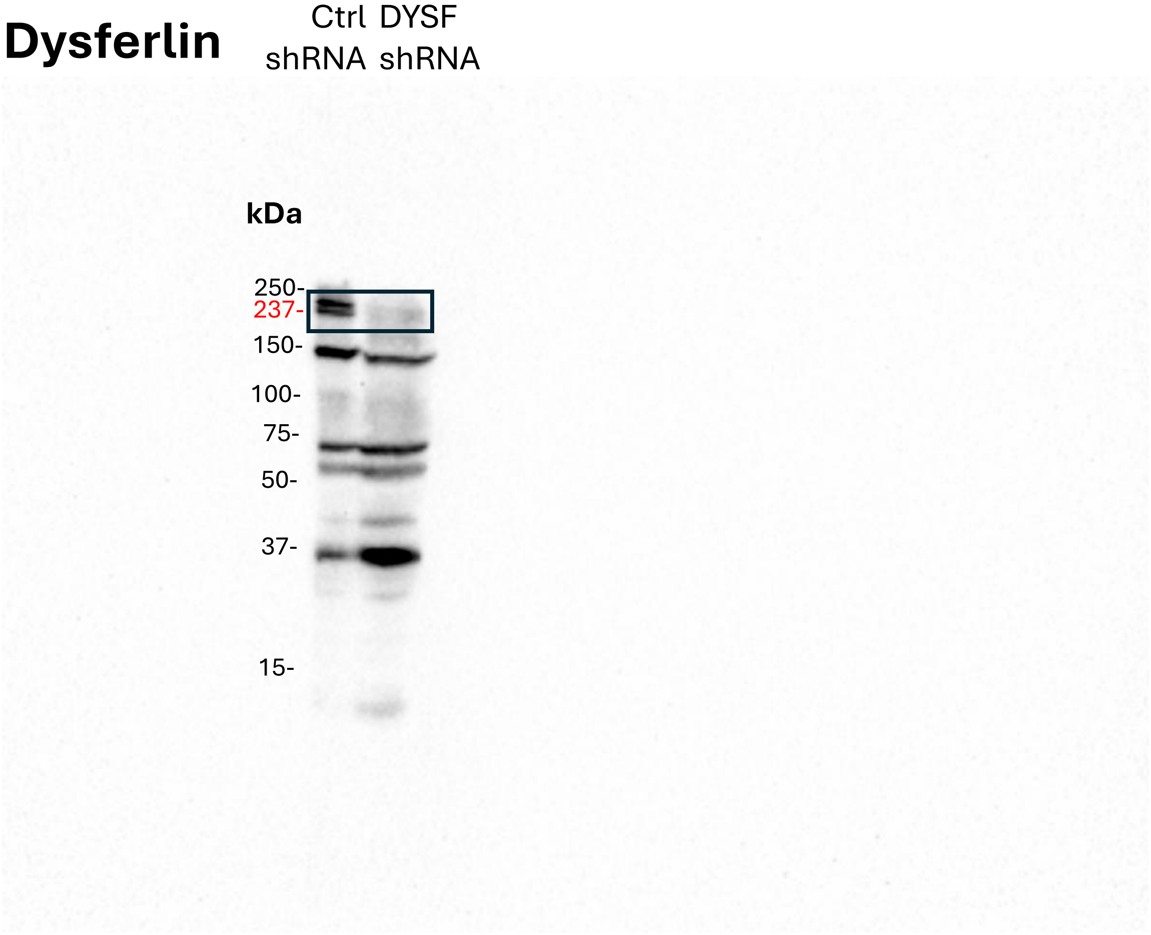

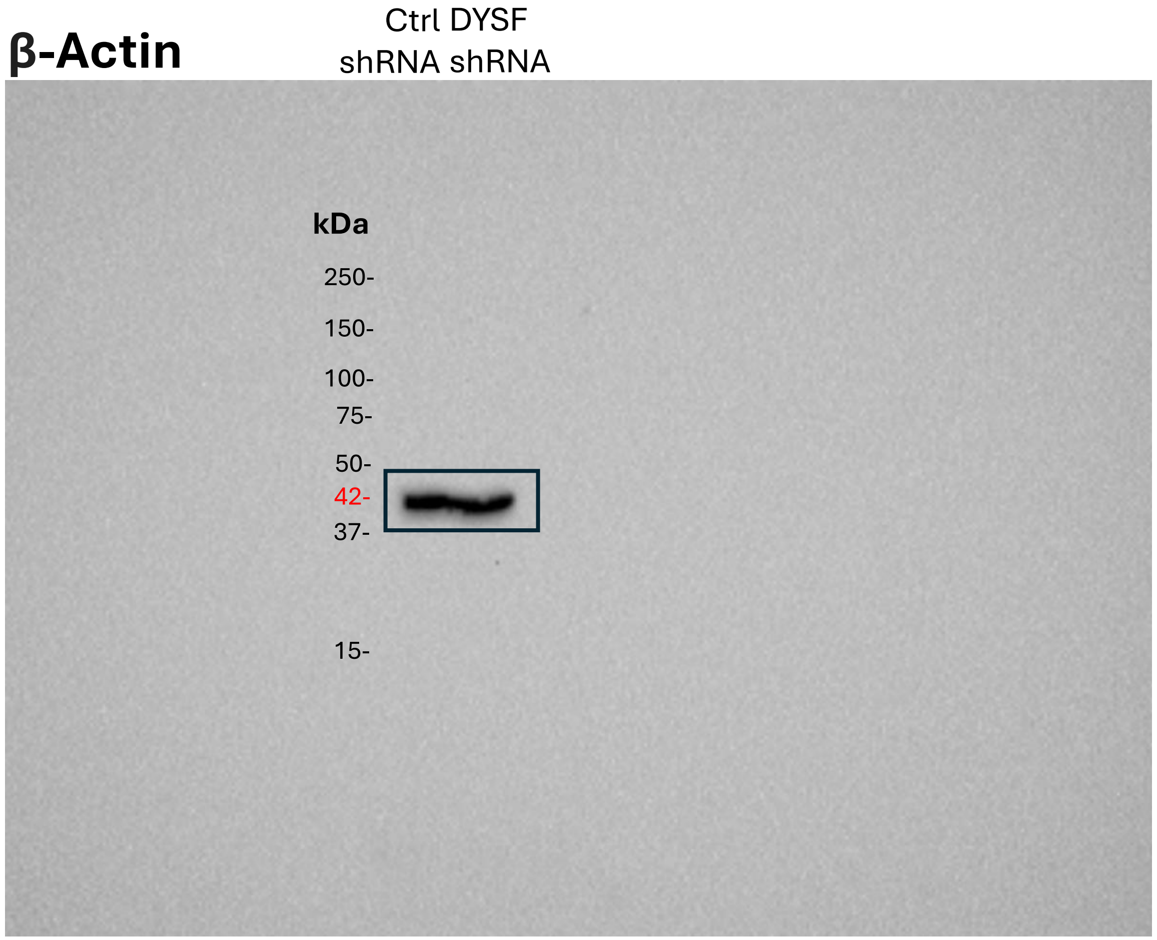

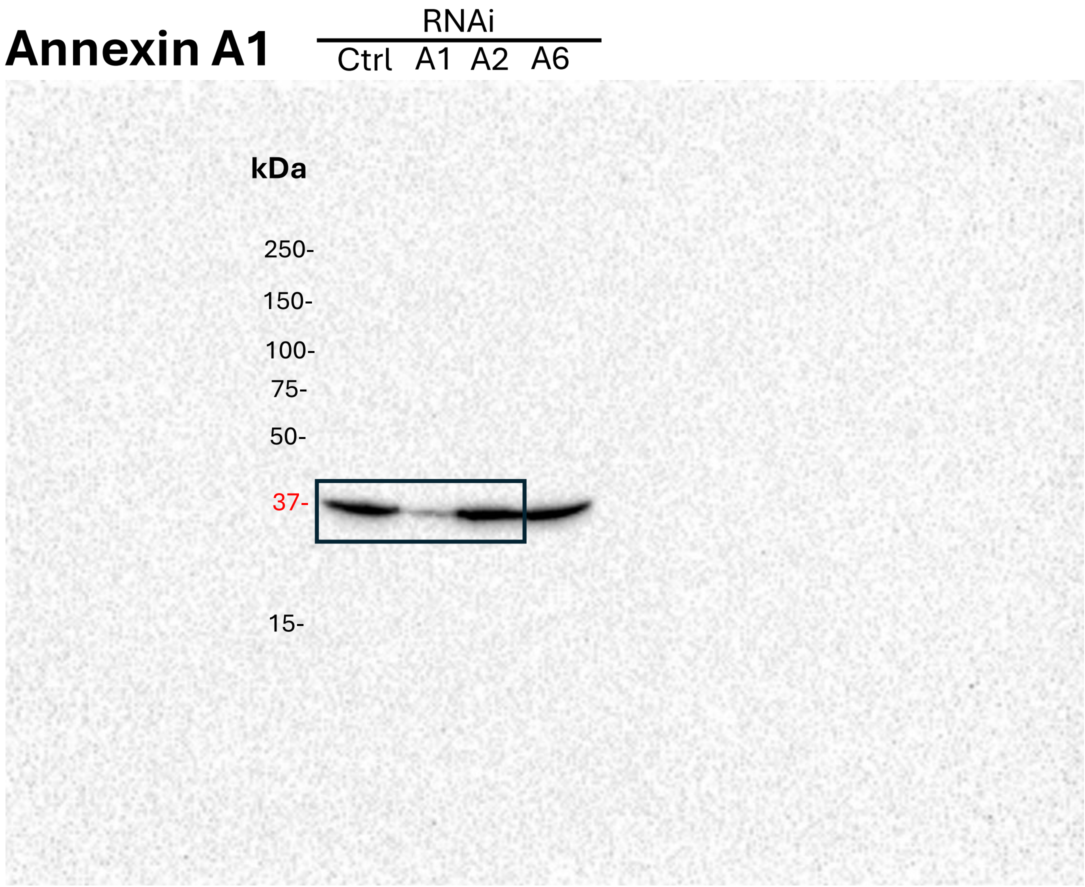

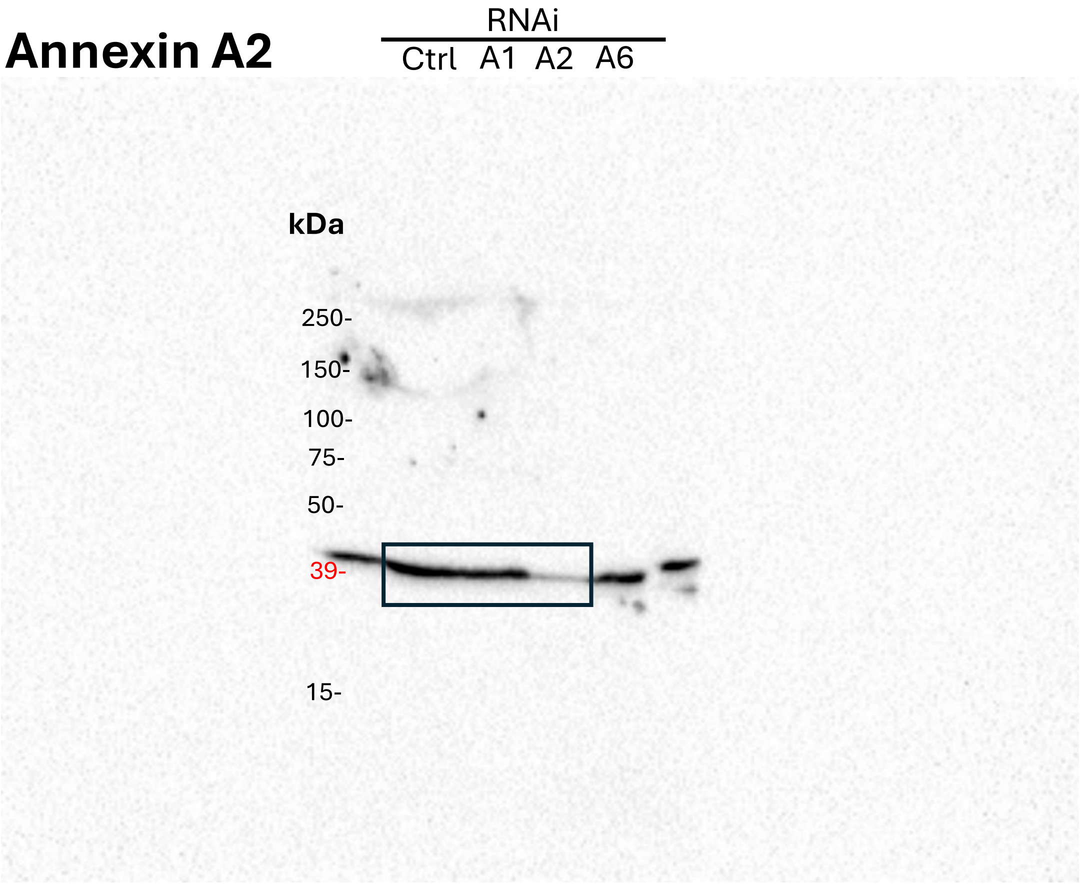

#### Figure 6C

##### β-Actin

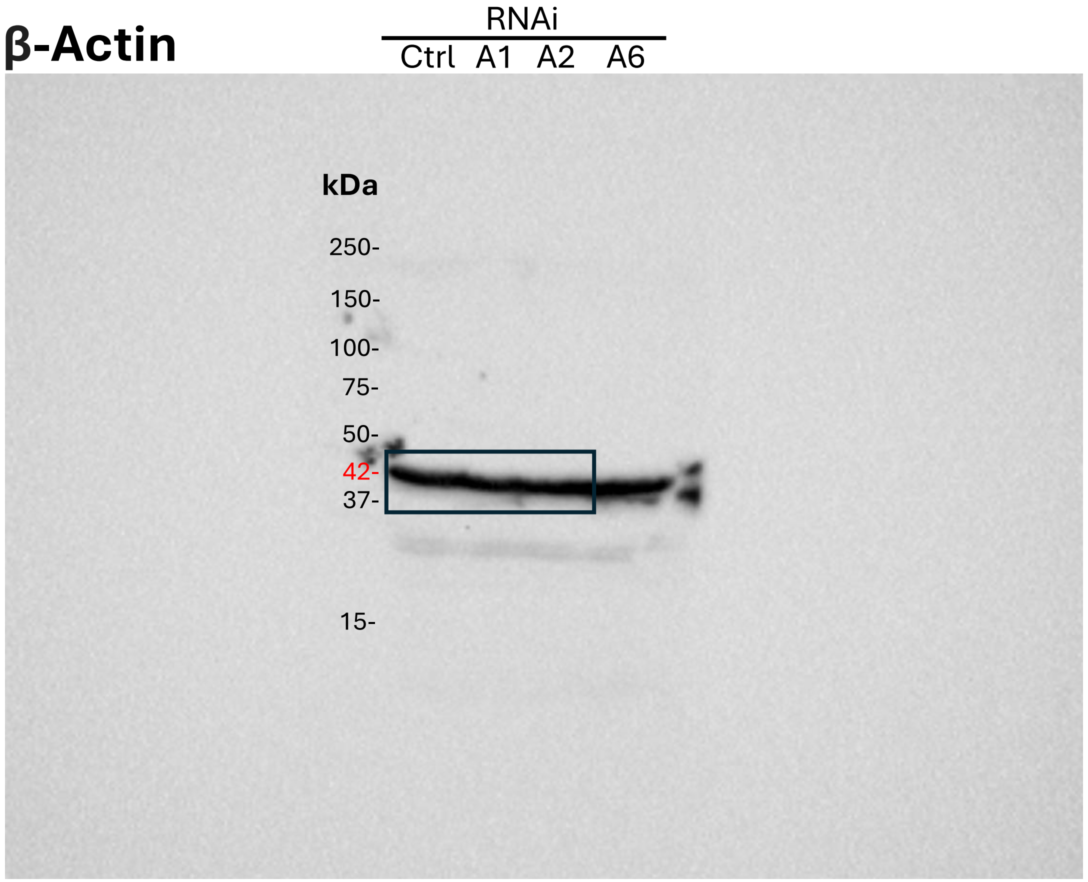

### Supplemental Fig S1

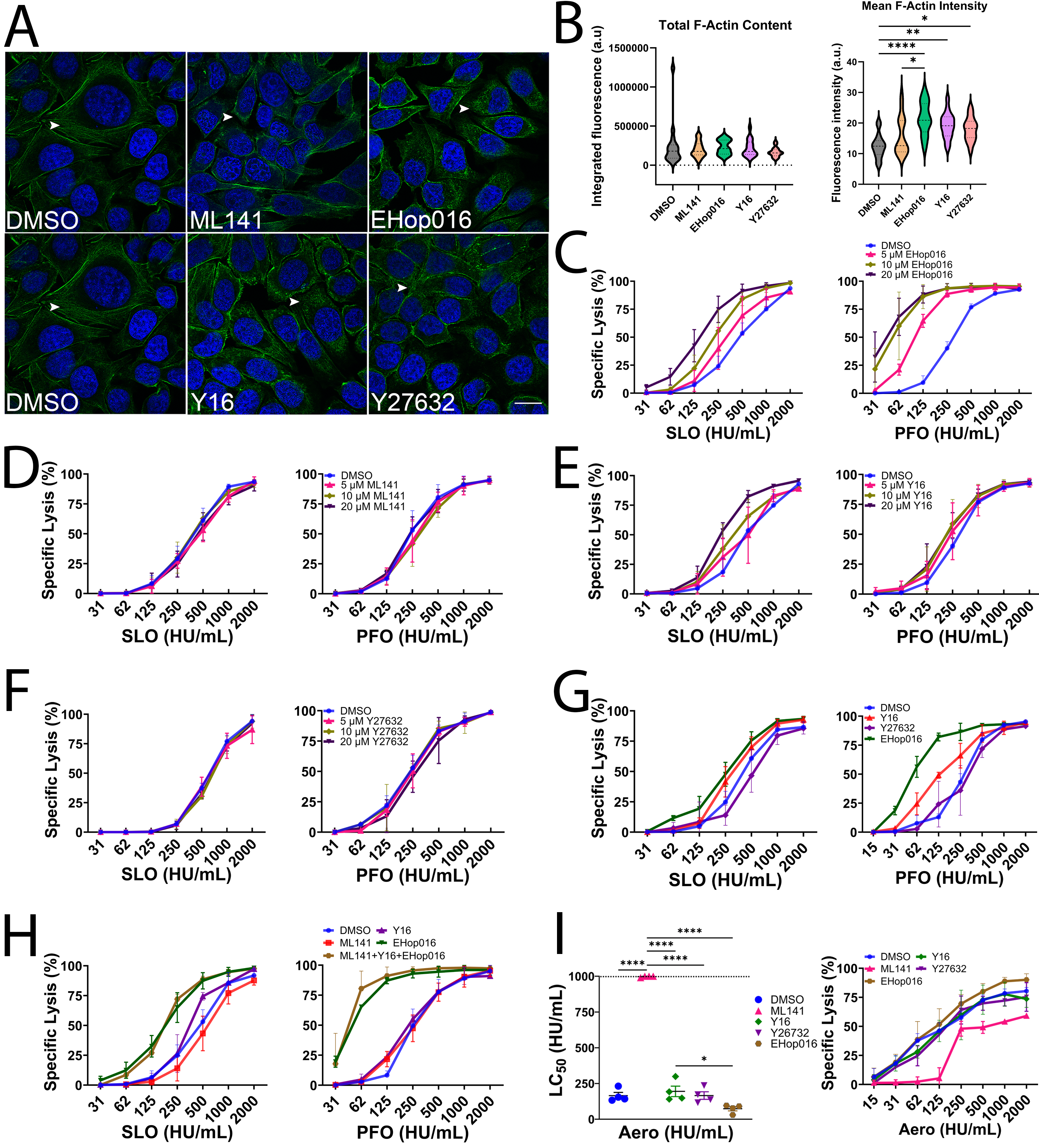

### Supplemental Fig S2

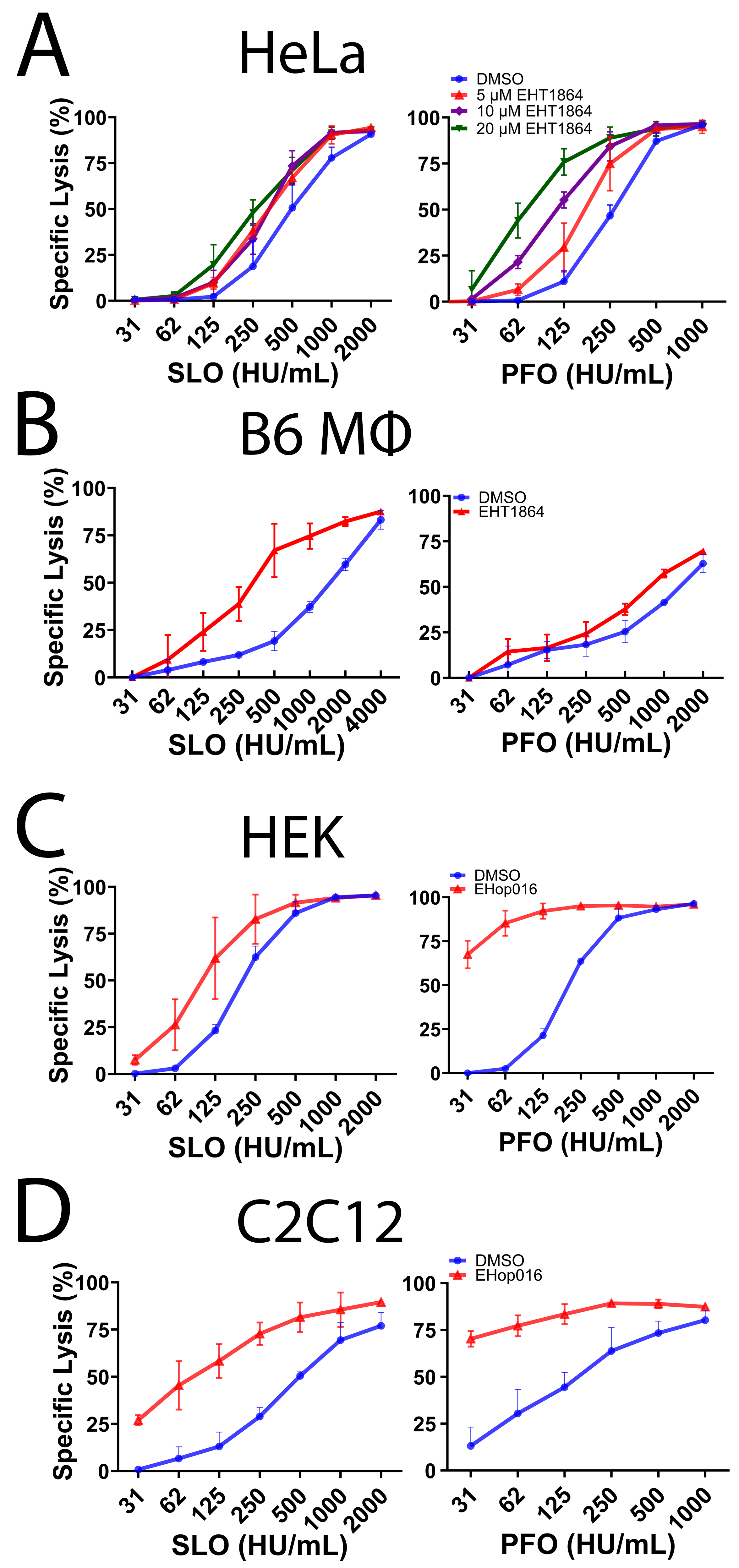

### Supplemental Fig S3

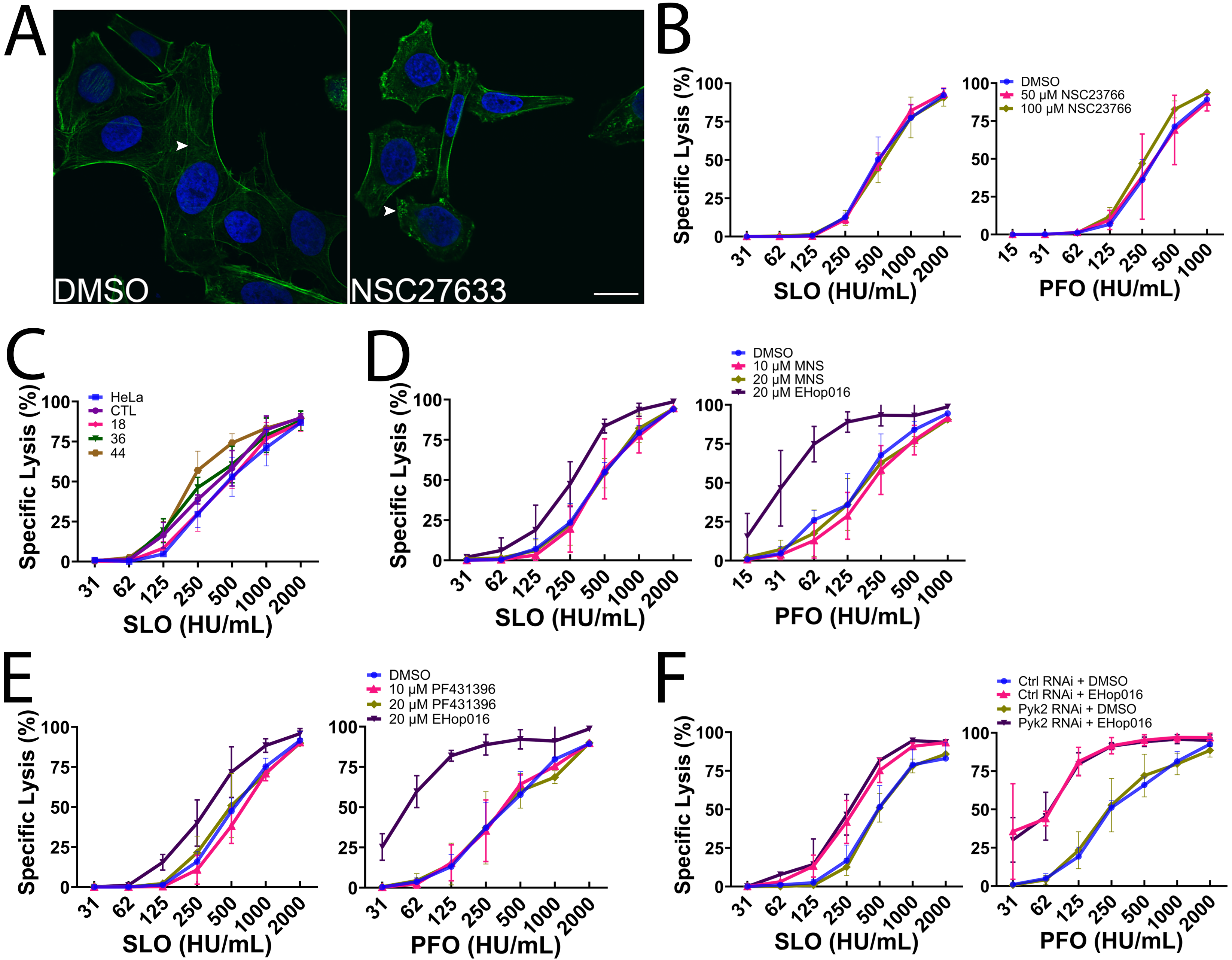

### Supplemental Fig S4

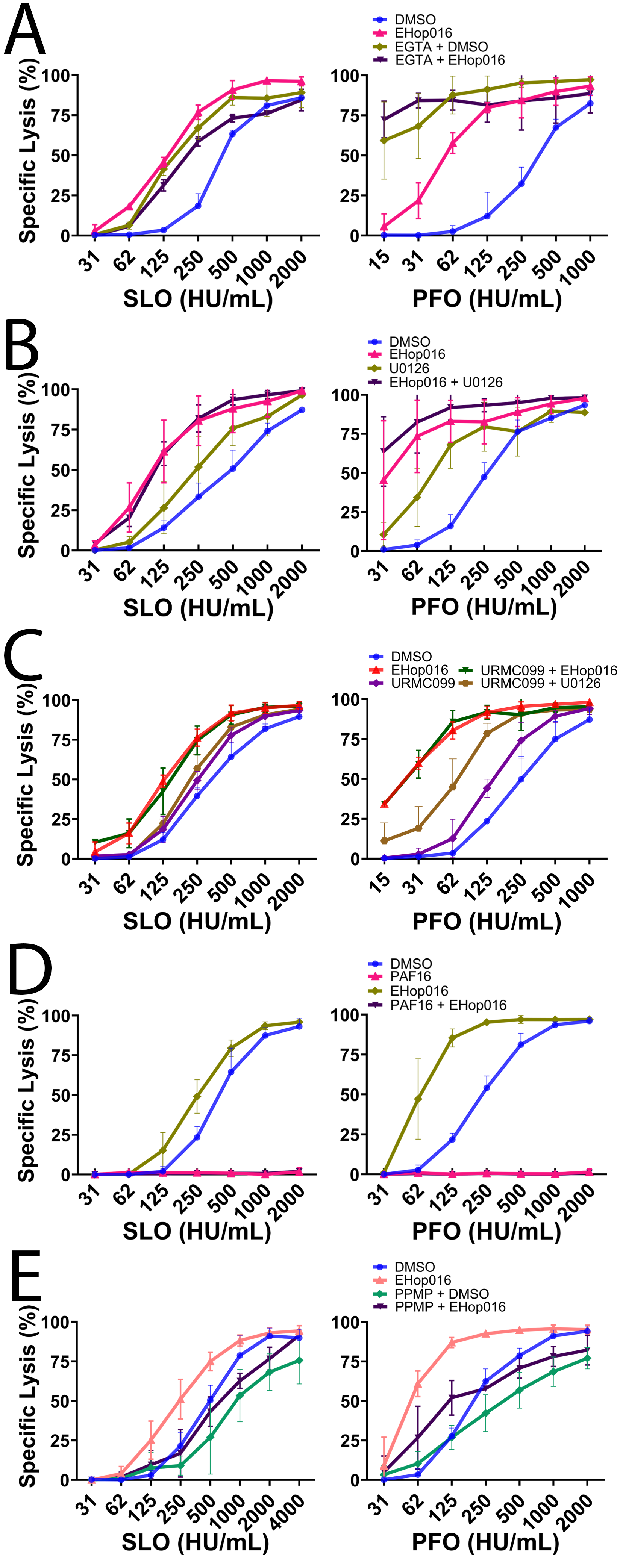

### Supplemental Fig S5

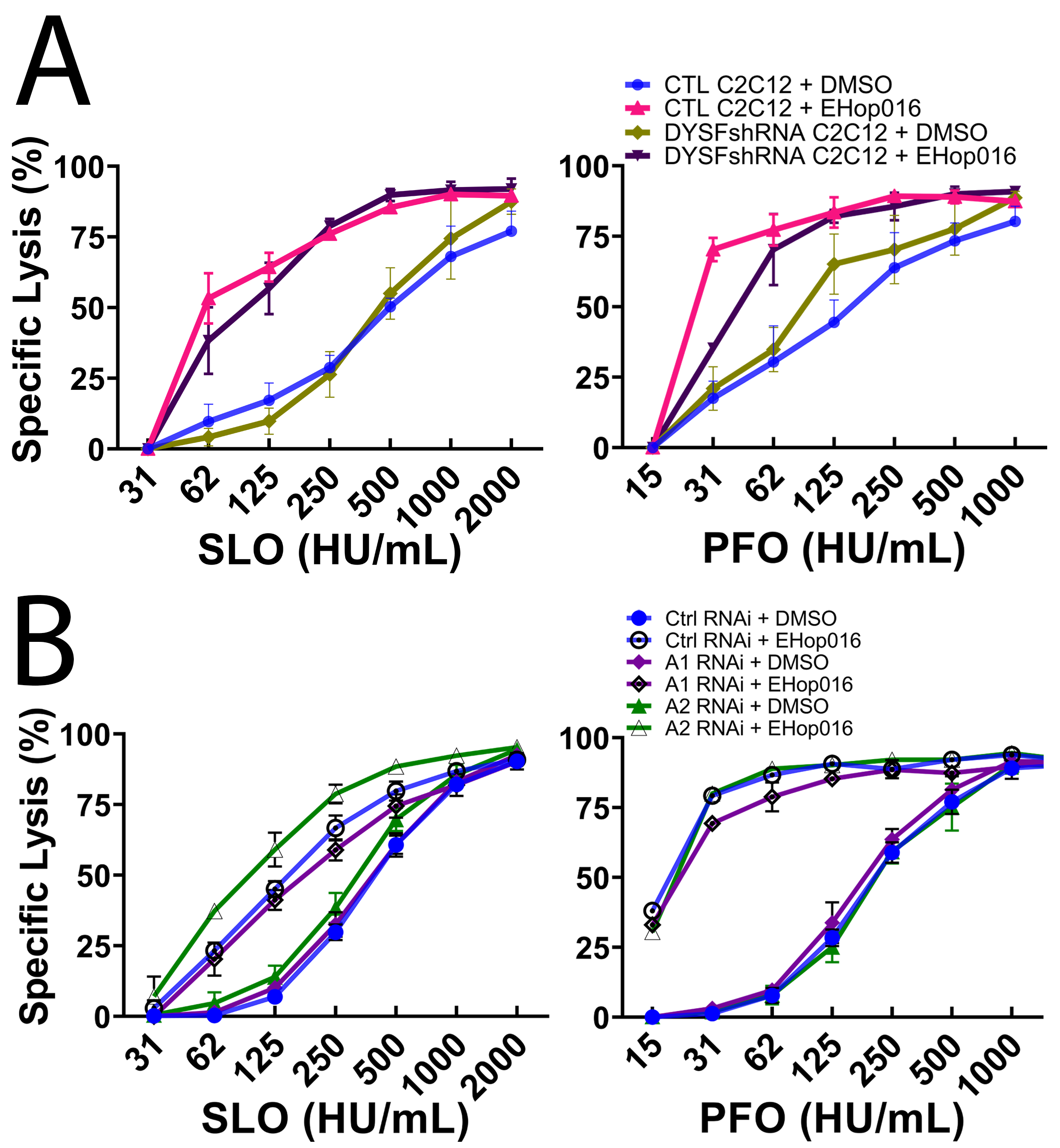
